# Dynamic trafficking and turnover of Jam-C is essential for endothelial cell migration

**DOI:** 10.1101/625913

**Authors:** Katja B. Kostelnik, Amy Barker, Christopher Schultz, Vinothini Rajeeve, Ian J. White, Michel Aurrand-Lions, Sussan Nourshargh, Pedro Cutillas, Thomas D. Nightingale

**Author notes:** These authors contributed equally (Katja B. Kostelnik, Amy Barker).

## Abstract

Junctional complexes between endothelial cells form a dynamic barrier that hinder passive diffusion of blood constituents into interstitial tissues. Re-modelling of junctions is an essential process during leukocyte trafficking, vascular permeability and angiogenesis. However, for many junctional proteins the mechanisms of junctional remodelling have yet to be determined. Here we used receptor mutagenesis, HRP and APEX-2 proximity labelling, alongside light and electron microscopy, to map the intracellular trafficking routes of junctional adhesion molecule-C (Jam-C). We found that Jam-C co-traffics with receptors associated with changes in permeability, such as VE-Cadherin, NRP-1 and 2, but not with junctional proteins associated with the transmigration of leukocytes. Dynamic Jam-C trafficking and degradation is necessary for junctional remodelling during cell migration and angiogenesis. By identifying new trafficking machinery we show that a key point of regulation is the ubiquitylation of Jam-C by the E3 ligase CBL, this controls the rate of recycling versus lysosomal degradation.

## Introduction

Endothelial cells exist as a monolayer that lines the blood vasculature and as such they present a discrete barrier to the cellular and molecular components of blood. This includes ions, small and large proteins, platelets and leukocytes^1^. The endothelium also plays a crucial role in responding to a pathogenic infection or tissue injury; both in the initial recruitment of leukocytes to the appropriate site^2^, and in subsequent tissue repair and angiogenesis^3^. Endothelial dysfunction can predispose the vessel wall to leukocyte adhesion, platelet activation, oxidative stress, thrombosis, coagulation and chronic inflammation leading to the pathogenesis of numerous cardiovascular diseases^4^.

Essential to the establishment of the endothelial monolayer are the protein complexes between adjacent cells that create junctions. Junctions are dynamic and are remodelled to control processes such as cell permeability^1^ and cell migration^5^, and a subset of junctional proteins also control leukocyte transmigration^2^. This subset includes molecules such as VE-Cadherin, PECAM-1 (CD31), CD99, endothelial cell-selective adhesion molecule (ESAM), ICAM-1 and -2 and the junctional adhesion molecule (Jam) family of proteins (Jam-A, -B and -C)^2, 6^. These proteins engage in homophilic and heterophilic interactions with neighbouring cells or with transmigrating leukocytes. Control of these interactions determines the stability of the junctions and/or provides unligated receptor for the movement of transmigrating leukocytes^2, 7^. For some receptors that are directly required for leukocyte transmigration (such as Jam-A and PECAM-1) this is a well characterised process regulated by a combination of phosphorylation, proteolytic cleavage and intracellular trafficking. However, for other receptors such as Jam-C almost nothing is known about the molecular control of receptor trafficking and its importance to function.

Jam-C is a type-1 integral membrane protein of the Ig superfamily that mediates numerous endothelial cell functions such as leukocyte trans-endothelial cell migration^8–12^, angiogenesis and vascular permeability^13–16^. However, there is a marked difference in the function of Jam-C compared to other endothelial junctional receptors. Rather than directly inhibiting leukocyte transmigration, inhibition of Jam-C by genetic means or using blocking antibodies changes the mode of transmigration^8, 11, 17, 18^. In Jam-C-deficient endothelial cells, leukocytes breach the endothelial barrier and then reverse, transmigrating back into the vessel lumen, reducing the overall efficiency of leukocyte traffic. Further, unlike Jam-A, Jam-C surface expression favours changes in permeability following stimulation with thrombin, VEGF or histamine by inhibiting the activation of the small GTPase Rap-1 and by activating actomyosin function^15^. Given these functional differences it is likely for the regulation of Jam-C to be similarly diverse. Together, the broad functional roles of Jam-C in inflammation and vascular biology is illustrated through its involvement in multiple inflammatory disease states such as arthritis^19^, peritonitis^9^, acute pancreatitis^18, 20^, ischemia reperfusion injury^10, 11^, pulmonary inflammation^21, 22^ and atherosclerosis^23–25^.

Whilst little is known about the molecular mechanisms of Jam-C trafficking, current evidence indicates an important role for such a phenomenon in regulating Jam-C function. Firstly, in cultured macro and microvascular endothelial cells Jam-C can be redistributed in a stimulus-dependent manner^13, 15, 26^; activation of endothelial cells with thrombin, VEGF or histamine increases localisation of Jam-C at the cell surface. Secondly, analysis of inflamed murine tissues by electron microscopy showed labelling of the extracellular domain of Jam-C within vesicles, at the cell surface and at junctional regions of endothelial cells. Importantly, distribution of Jam-C was altered within these sites following an inflammatory stimulus (ischemia/reperfusion injury), resulting in a change in the levels of vesicular protein vs. junctional and cell surface protein^10, 11^. Finally, enhanced levels of Jam-C have been observed in atherosclerosis and rheumatoid arthritis, some of which likely reflect changes in intracellular trafficking^19, 24^.

Collectively current evidence suggests a causal link between redistribution of intracellular Jam-C and physiological and pathological processes involving opening of endothelial cell junctions. To directly address this hypothesis, we investigated the mechanisms underlying Jam-C redistribution and trafficking using wild-type and mutant variants of this receptor and a combination of light and electron microscopy. Furthermore, to identify co-trafficked cell surface receptors and trafficking machinery involved in Jam-C internalisation and subcellular localisation, we developed novel proximity labelling mass spectrometry approaches. By perturbing the newly identified pathways of Jam-C re-distribution, and using mutant variants of this receptors, we show that a key point of regulation is the ubiquitylation of Jam-C by the E3 ligase CBL, this regulates the rate of recycling versus lysosomal degradation.

## Methods

### Constructs and cloning

pRK5-HA-Ubiquitin-WT was a gift from Ted Dawson (Addgene plasmid # 17608). Murine Jam-C-GFPout and Jam-C-GFPout ΔPDZ have been described previously^27^. To generate Murine Jam-C-HRPout, HRP was amplified from P-selectin HRP^28^ using primers incorporating 15 bp 5’ or 3’ overlaps with Jam-C (Supp. Table 1). Jam-C-GFPout was digested with *Xba*I and *Xho*I and an in-fusion reaction and subsequent transformation performed according to the manufacturer’s instructions (Clontech, Mountain View, CA). QuikChange site directed mutagenesis of Jam-C-GFPout to generate Y267A, S281A, Y282A, K283R, K287R, Y293A and T296A was performed according to the manufacturer’s instructions (Agilent, Santa Clara, CA) using the primers listed (Supp. Table 1). Murine Quad-K Jam-C-HRPout, Jam-C-APEX2in and all the site directed mutations thereof were prepared by gene synthesis (Thermofisher, Waltham, MA). The soy bean sequence was first codon optimised using GeneOptimizer (Thermofisher) and incorporated into the intraluminal domain of Jam-C between the transmembrane and cytoplasmic tail before subcloning into pCDNA3.1 using *Hind*III and *Bam*HI sites. For lentiviral transduction WT and Quad-K Jam-C-GFPout were amplified using appropriate primers (Supp. Table 1) and subcloned into pSFFV-WPRE using an infusion reaction according to the manufacturer’s instructions (Clontech).

### Cell Culture and transient transfection

Human Umbilical Vein Endothelial Cells (HUVEC, Promocell, Heidelberg, Germany) were cultured as previously described^29^. Plasmid transfections were performed by nucleofection (Nucleofector II, programme U-001, Amaxa Biosystems, Gaithersburg, MD) using 2-10 μg DNA or 250 pmol siRNA (Jam-C-AUGUAGUUAACUCCAUCUGGUUUCC or CBL-1-CCUCUCUUCCAAGCACUGA or CBL-2-CCUGAUCUGACUGGCUUAU).

### Lentivirus preparation

To produce lentivirus, 8×10^6^ HEK293T cells were seeded onto a 15 cm diameter tissue culture dish in 20ml DMEM with Glutamax (Gibco) with 10% heat inactivated FBS (Gibco). Cells were incubated overnight before adding 25 μM chloroquine solution 1 hour prior to transfection. Transfection complexes were prepared by diluting 18 μg of plasmid expressing the lentiviral packaging genes (pGagPol), 4 μg plasmid expressing the VSV-G envelope (pMDVSVG), and 18 μg of pSFFV-JAM-C-EGFP-WPRE in 2ml OptiMEM (Gibco). 200μl polyethylenimine (PEI) solution (1mg/ml) was added (5:1 ratio of PEI:DNA) and immediately pulse vortexed (3 brief pulses, low speed so as not to shear DNA). Complexes were incubated at room temperature for 10 minutes before all volume was transferred to HEK293T cells, which were returned to the incubator overnight before media was aspirated and replaced with 20 ml fresh DMEM for 48 hours. Virus-containing supernatant was removed and stored at 4°C, and replaced with another 20 ml DMEM complete for a further 24 hours. The supernatant was taken and pooled with the earlier material, syringe filtered with a 0.22 μM pore filter. 4X concentrated polyethylene glycol (PEG) precipitation solution was prepared (36% PEG (6,000 Da), 1.6 M NaCl, filter sterilised), and diluted to 1X concentrate in the virus containing supernatant. This was stored at 4°C for 90 minutes (inverted every 30 min) before centrifugation at 1500 x g for 1 hour at 4°C, following which media was aspirated and the pellet resuspended in OptiMEM. Aliquots were stored at −80°C until use.

### Antibody feeding

HUVEC were cultured on 10 mm coverslips and inverted on pre-warmed drops of HUVEC growth medium (HGM) containing 1:1000 dilution of Rabbit anti-Jam-C antibody (Supp. Table 2) for 15 min. Coverslips were then washed 3 times in pre-warmed PBS and transferred to a drop of HGM. Coverslips were fixed at 0, 15, 30 min, 1, 2 or 4 h and immunofluorescence labelled for total or cell surface antibody and the nuclei labelled with DAPI before analysis by confocal microscopy. Levels of cell surface Jam-C antibody were determined using Cell Profiler software™^30^. Nuclei and edge channels were manually Otsu thresholded and a median filter was applied with an artificial diameter of 10 pixels. Objects less than 50 pixels in diameter were filtered out and the object area and intensity determined.

### Ubiquitylation assay

Endogenous CBL was depleted in HUVEC by two rounds of transfection with 250 pmol siRNA as above. At the second round of transfection pRK5-HA-Ubiquitin-WT and WT or mutant Jam-C-GFPout were included, the next day cells were lysed in RIPA buffer (150 mM NaCl, 1% NP40, 0.5% sodium deoxycholate, 0.1% SDS and 50mM Tris pH 8.0) supplemented with 10 mM freshly prepared N-ethylmaleimide and protease inhibitors (Sigma-Aldrich). Washed GFP-Trap A beads (Chromotek) were added to the lysates and incubated overnight rotating at 4°C. Beads were washed and resuspended in hot laemmli buffer (2% SDS, 25% glycerol, 0.36 M β-mercaptoethanol, 0.05 M Tris pH8.0) and western blotted.

### HRP and APEX-2 proteomics

Typically, 4X 14 cm plates (Nunc, Roskilde, Denmark) of HUVEC were required for each proteomics condition analysed. Endogenous Jam-C was depleted by two rounds of transfection over a 96 h period with 250 pmol Jam-C siRNA/reaction. At the second round 10 μg WT, QuadK Jam-C-HRPout or APEX-2in constructs were also included. Following the second round transfection cells were cultured with 7 μM freshly made heme to aid peroxidase folding and in the presence or absence of 50 ng/ml TNFα and/or 100 nM Bafilomycin (Life Technologies). 24 h after the second transfection, cells were fed with 500 μM biotin tyramide (Iris Biotech, Marktredwitz, Germany) for 30 min at 37°C. The cells were then exposed to M199 supplemented with 1 mM Hydrogen Peroxide in the presence or absence of 50 mM freshly prepared ascorbate for 1 min. The biotinylation reaction was stopped by the addition of stop solution (PBS, 10 mM sodium azide, 10 mM ascorbate, 5 mM Trolox). Cells were either fixed for immunofluorescence analysis or lysed at 4°C in RIPA buffer supplemented with 10 mM sodium azide and protease inhibitors. The lysate was centrifuged at (21000 g) for 15 min at 4°C and protein concentration determined (Pierce^™^ 660nm Protein Assay Reagent, Thermo Scientific). 8.5 μg lysate was kept for western blot analysis and 1.6-1.8 mg total protein was added to 250 μl washed high capacity neutravidin beads (Life Technologies) in low binding tubes (Life Technologies) and rotated overnight at 4°C. The beads were washed in 25 mM ammonium bicarbonate buffer, centrifuged at 21000 x g and frozen at −80°C before mass spectrometry analysis.

### Cycloheximide chase assay

HUVEC were transfected with WT or mutant Jam-C-GFPout, the next day cells were washed and incubated with 10 μg/ml cycloheximide. Cells were lysed in RIPA buffer supplemented with protease inhibitors (Sigma-Aldrich) at 0, 8 and 24 h before analysis by western blotting. Mass Spectrometry

Proteomics experiments were performed using mass spectrometry as reported^31, 32^. In brief, Immunoprecipitated (IP) protein complex beads were digested into peptides using trypsin and peptides were desalted using C18+carbon top tips (Glygen corporation, TT2MC18.96) and eluted with 70% acetonitrile (ACN) with 0.1% formic acid. Dried peptides were dissolved in 0.1% TFA and analysed by nanoflow ultimate 3000 RSL nano instrument coupled on-line to a Q Exactive plus mass spectrometer (Thermo Fisher Scientific). Gradient elution was from 3% to 35% buffer B in 120 min at a flow rate 250nL/min with buffer A being used to balance the mobile phase (buffer A was 0.1% formic acid in water and B was 0.1% formic acid in ACN).

The mass spectrometer was controlled by Xcalibur software (version 4.0) and operated in the positive mode. The spray voltage was 1.95 kV and the capillary temperature was set to 255 ºC. The Q-Exactive plus was operated in data dependent mode with one survey MS scan followed by 15 MS/MS scans. The full scans were acquired in the mass analyser at 375-1500m/z with the resolution of 70 000, and the MS/MS scans were obtained with a resolution of 17 500. The mass spectrometry proteomics data have been deposited to the ProteomeXchange Consortium via the PRIDE partner repository with the dataset identifier PXD013003. MS raw files were converted into Mascot Generic Format using Mascot Distiller (version 2.5.1) and searched against the SwissProt database (release December 2015) restricted to human entries using the Mascot search daemon (version 2.5.0) with an FDR of ~1%. Allowed mass windows were 10 ppm and 25 mmu for parent and fragment mass to charge values, respectively. Variable modifications included in searches were oxidation of methionine, pyro-glu (N-term) and phosphorylation of serine, threonine and tyrosine. The mascot result (DAT) files were extracted into excel files for further normalisation and statistical analysis.

### Calcium switch

HUVEC were washed in low calcium media (M199 with 20% dialysed FBS-SigmaAldrich and 10 U/ml Heparin) before addition of low calcium media with 4 mM EGTA pH 8.0. After 30 min the EGTA was removed and replaced with HUVEC growth media. Cells were fixed before and 30 min after EGTA addition and at 15, 30, 60 and 90 min post washout. Cells were then analysed by confocal microscopy labelling for Jam-C and VE-cadherin.

### Live cell Imaging

Endogenous Jam-C or CBL was depleted in HUVEC by two rounds of transfection with 250 pmol siRNA as above. At the second round cells were transfected with WT or Quad K Jam-C-GFPout and plated on borosilicate glass bottomed dishes (Greiner Bio One, Kremsmunster, Austria). The following day cells were imaged in the presence or absence of 100 nM Bafilomycin (Life technology) in a heat-controlled chamber at 37°C with 5% CO_2_ in HUVEC growth media either using 63x oil immersion objective (NA 1.3) and a Zeiss 800 microscope (Zeiss, Jena, Germany) or for higher speed images using a 100x oil immersion lens (NA 1.4) and a spinning disk (UltraVIEW VoX; Perkin-Elmer). During longer term timelapse (45 min) images were acquired every 45 s at a resolution of 512 x 512 pixels and a step size of 0.5 μm. During spinning disk acquisition images were acquired every 5 s for 10 min with a step size of 0.4–0.5 μm, comprising 9–14 pictures (depending on cell height) with an exposure for each image at 30 ms.

### Immunofluorescence staining

Fixation and staining were carried out as in Lui-Roberts et al.^33^ using appropriate antibodies (see Supp. Table 2). Fixed cell images were taken on a Zeiss 800 scanning confocal microscope system with a 63x objective (NA 1.3) as confocal z-stacks with 0.5 μm step size. Acquisition was performed using Zen Blue software with a 1024×1024 pixel resolution, 2x frame average and 1x zoom.

### Western blotting

Proteins were separated by SDS-PAGE, transferred to Polyvinylidene fluoride membranes (PerkinElmer), and then probed with primary antibody (see Supp. Table 2) followed by the appropriate HRP-conjugated secondary antibody (1:5000) (Agilent, Santa Clara, CA).

### Scratch wound assay

Mock or Jam-C knock down cells were transduced with GFP, WT Jam-C-GFPout or QuadK Jam-C-GFPout lentivirus. Scratch wound migration assays were performed on confluent monolayers of HUVEC. Wounds were made using a pipette tip and images were captured every 30 min for 16 h by time lapse microscopy using an Olympus (Shinjukum, Tokyo) IX81 microscope, additional images of closed scratch wounds were also acquired at 60 h. Percentage wound closure was calculated using Fiji ^34^.

### Electron microscopy

HUVECs were transfected with WT or Quad-K Jam-C-HRPout constructs and plated onto glass coverslips. The coverslips were fixed in EM-grade 2% paraformaldehyde, 1.5% glutaraldehyde (TAAB Laboratories Equipment, Ltd., Aldermaston, UK) and washed in 0.05M Tris HCl (pH7.6) before incubation with 0.075% diaminobenzidine (DAB); 0.02% hydrogen peroxide for 30 min in the dark. The coverslips were washed in 0.1M sodium cacodylate and secondarily fixed in 1% osmium tetraoxide; 1.5% potassium ferricyanide and then 1% tannic acid treated. Samples were then dehydrated and embedded in Epon resin. Coverslips were inverted onto pre-polymerized Epon stubs and polymerized by baking at 60°C overnight. 70 nm-thin sections were cut with a diatome 45° diamond knife using an ultramicrotome (UC7; Leica, Wetzlar, Germany). Sections were collected on 1 × 2 mm Formvar-coated slot grids and stained with Reynolds lead citrate. Samples were imaged using a transmission electron microscope (Tecnai G2 Spirit; FEI, Thermofisher) and a charge-coupled device camera (SIS Morada; Olympus).

## Results

### Jam-C is dynamically trafficked from the cell surface

To characterise the amount and localisation of intracellular Jam-C present at steady state in cultured endothelial cells, we carried out immunofluorescence analysis in human umbilical vein endothelial cells (HUVEC) (Fig. 1A-D). In confluent monolayers, Jam-C is primarily localised to the junctions although punctae of Jam-C are also present in just over half of the cells (Fig. 1D). Incubating cells with the vacuolar-type H+-ATPase inhibitor bafilomycin (100nM for 4 h) (Fig. 1C), a reagent that blocks the acidification step essential for lysosomal degradation, markedly increased the number of Jam-C-positive intracellular punctae. These results indicate that Jam-C is constitutively trafficked into vesicles, and at least a proportion of the receptor in resting conditions is targeted for degradation in lysosomes.

**Figure. 1.**
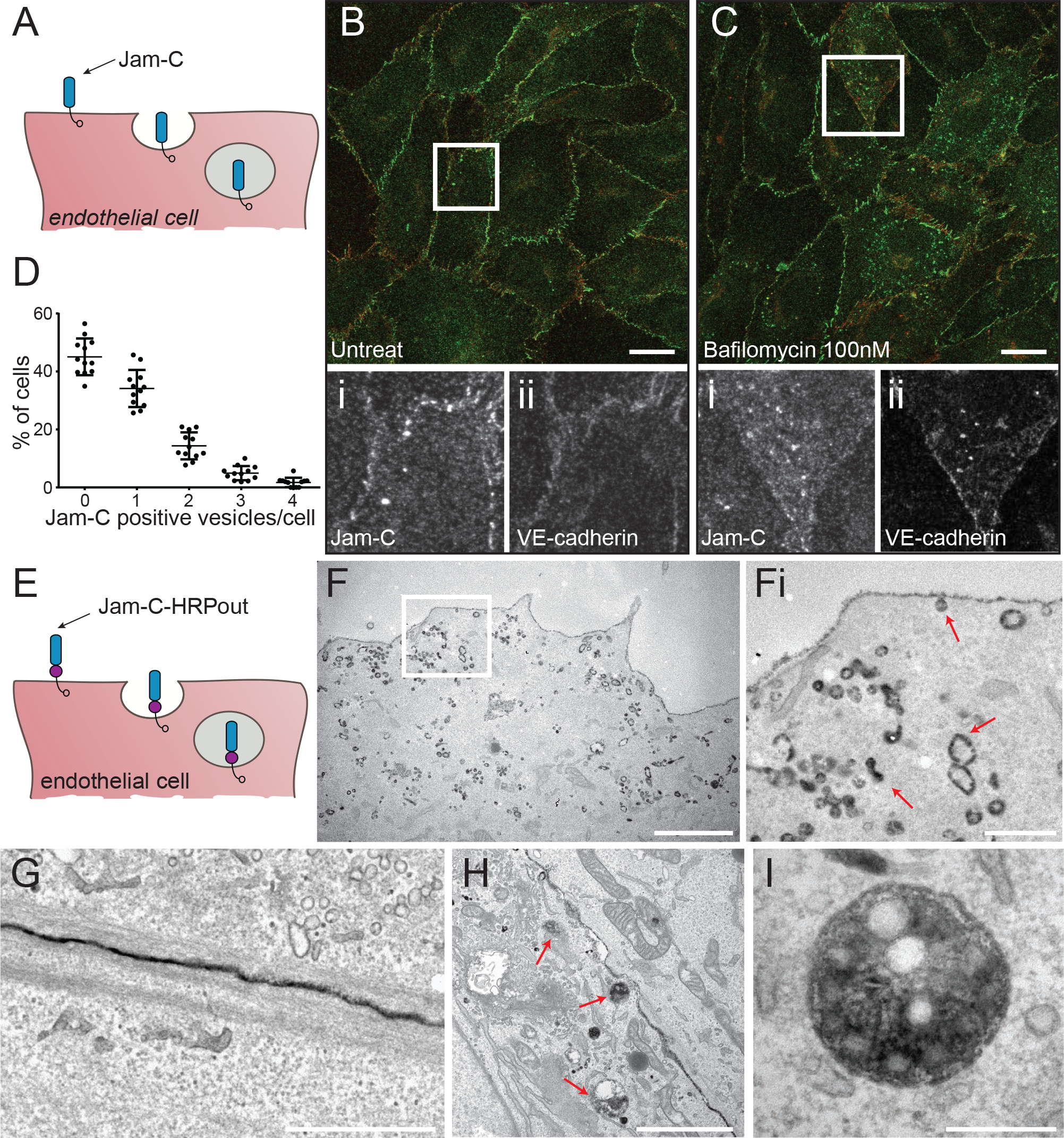
Jam-C is constitutively trafficked from endothelial cell junctions. (A) Schematic of the endocytosis of endogenous Jam-C from the endothelial cell surface showing the extracellular immunoglobulin domains (blue) and the intracellular PDZ interacting domain (clear circle). (B) Untreated or (C) 100nM bafilomycin treated HUVEC were fixed and co-stained for Jam-C (green) and VE-Cadherin (red). Boxed regions are shown magnified below in greyscale (i) Jam-C (ii) VE-Cadherin. Scale bar 20μm. (D) Quantification of the % of untreated cells with a number of Jam-C positive vesicles (551 cells from n=3 experiments, error bars represent SEM). (E) Schematic of the endocytosis of Jam-C-HRPout from the endothelial cell surface showing the extracellular immunoglobulin domains (blue), the HRP tag (purple circle) and the intracellular PDZ interacting domain (clear circle). (F-I) HUVEC were transiently transfected with Jam-C-HRPout, fixed and incubated with diaminobenzidine (DAB) and hydrogen peroxide for 30min. Cells were then secondarily fixed and 70nm sections prepared and imaged by transmission electron microscopy. Precipitated DAB can be clearly seen on (F) endocytic structures and early endosomes, (Fi) shows boxed area magnified, red arrows highlight endocytic structures and endosomes, (G) junctions, (H and I) multi-vesicular bodies (red arrows). Scale bars (F) 2 μm (Fi) 500 nm (G) 1 μm (H) 2 μm (I) 200 nm.

We next determined the ultrastructure of intracellular Jam-C pools using electron microscopy (EM). We generated a version of Jam-C fused to horse radish peroxidase (HRP) in the membrane proximal region of the extracellular domain (Jam-C-HRPout). Tagging of Jam-C at this point has previously been shown to have no effect on ligand binding or localisation^27^. We utilised murine Jam-C for expression studies in HUVEC as this allows the depletion of endogenous human Jam-C and rescue with the siRNA-resistant murine orthologue. Cells expressing Jam-C-HRPout were fixed and labelled with diaminobenzidine (DAB) before analysis by electron microscopy. DAB labelling was present at the junctions (Fig. 1G), in small endocytic clathrin-negative structures budding off from the cell surface (Fig. 1F & 1Fi), endosomes (Fig. 1F) and in late endosomes/multi-vesicular bodies (MVBs) (Fig. 1H & I).

To more accurately determine the kinetics of endogenous Jam-C internalisation, we fed anti-Jam-C antibody and monitored the fluorescent intensity at the junctions over a 2 h period (Fig. 2A & B). This approach showed that approximately 2/3rds of surface protein are removed from the junction within 2 h (Fig. 2B) indicating that Jam-C is rapidly removed from the junctions at steady state.

**Figure. 2.**
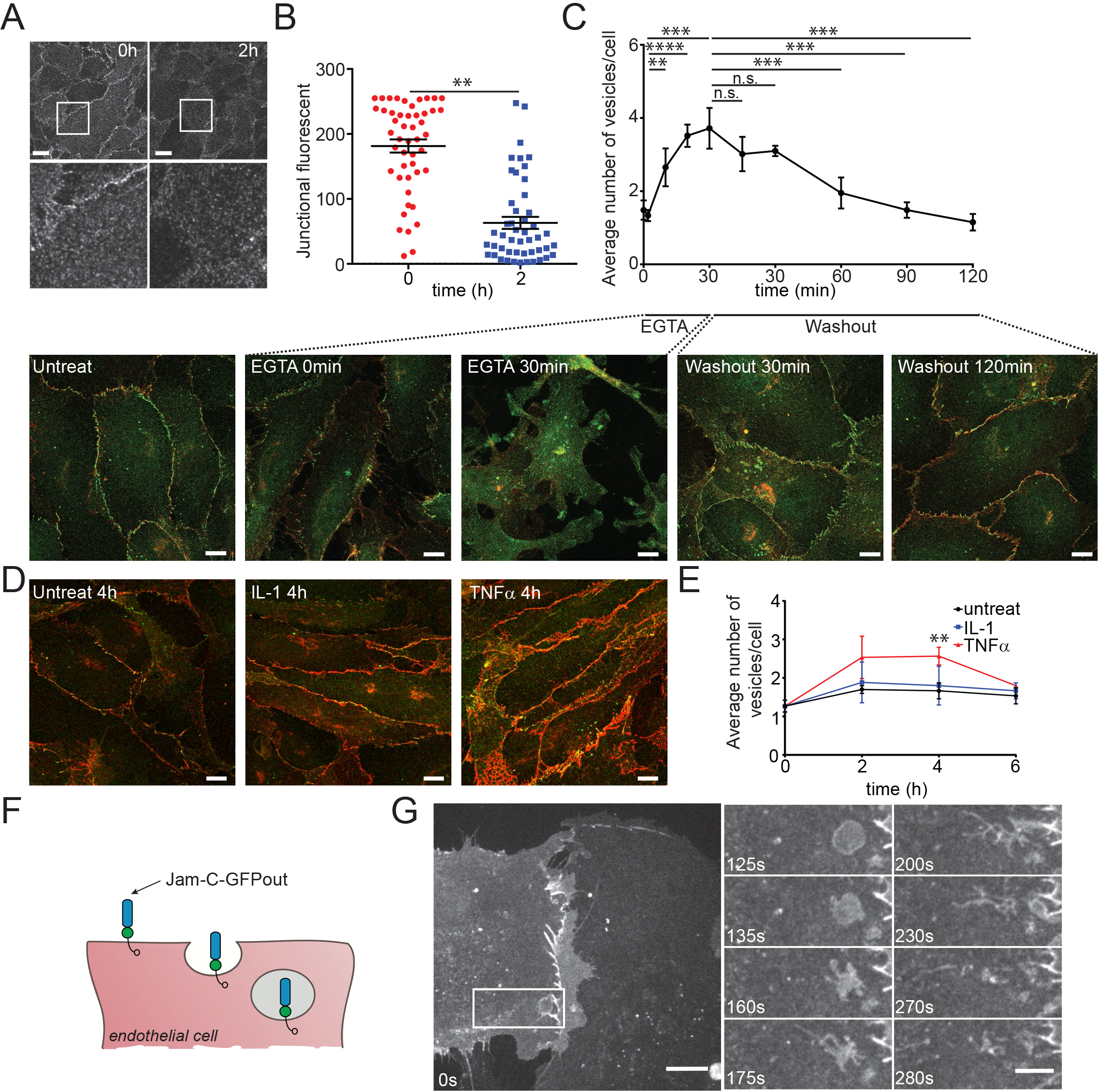
Jam-C is turned over dynamically at the endothelial junction and the rate of internalisation is increased during junctional disassembly and inflammation. (A) HUVEC were fed with Jam-C antibody, excess antibody was removed and the cells were incubated at 37°C before being fixed at 0 and 2 h and imaged by confocal microscopy. Scale bar 20 μm, boxed regions are shown at higher magnification. (B) The difference in junctional intensity is quantified between 0 and 2 h (n=4 experiments, error bars represent SEM, **P≤0.01; t-test). (C) HUVEC were treated with 4 mM EGTA in calcium free low serum media for 30 min before washing and incubation at 37°C with conventional media. Cells were fixed before and during the time course of wash out and recovery and stained for Jam-C (green) and VE-Cadherin (red). Images were acquired by confocal microscopy and the number of Jam-C positive vesicles at each time point determined. Scale bar 10 μm (5 fields of view/experiment, n=3 experiments, error bars represent SD, **P ≤0.01, ***P≤0.001, ****P≤0.0001; t test). (D) HUVEC were either left untreated or incubated with 10 ng/ml IL-1 or 50 ng/ml TNFα for 4h at 37°C. Cells were then fixed and stained for Jam-C (green) and VE-Cadherin (red). Images were acquired by confocal microscopy and (E) the number of Jam-C positive vesicles at each time point determined. Scale bar 10 μm (150 cells from n=3 experiments, error bars represent SD, **P≤0.01; t test). (F) Schematic of the endocytosis of Jam-C-GFPout from the endothelial cell surface showing the extracellular immunoglobulin domains (blue), the GFP tag (green circle) and the intracellular PDZ interacting domain (clear circle). (G) HUVEC were nucleofected with WT JAM-C-GFPout and imaged with a spinning-disk confocal microscope. Time indicates total time in media and the boxed region at 0 s is shown magnified at later time points. Jam-C exists in vesicles, at the cell surface and at the cell junctions. Large vesicles can be seen that tubulate and migrate away from the junctional region. Scale bar: 0s, 20 μm; inset, 10 μm.

Endothelial cell junctions are remodelled during vascular growth, angiogenesis and inflammation, allowing cells to move within the monolayer and leukocytes to cross the endothelial barrier. Once these processes are complete it is essential for the junctions to reform to once again establish a contiguous barrier. Using immunofluorescence and confocal microscopy to analyse localisation of endogenous Jam-C, we monitored the amount of vesicular traffic associated with conditions that are known to disrupt endothelial barrier function. We first artificially disrupted junctions by chelating calcium ions essential for junctional formation^35^; we then washed out the chelating agent (EGTA) to monitor junctional reformation (Fig. 2C). An increased number of Jam-C positive vesicles was noted following the addition of EGTA that then reduced following washout and formation of a confluent monolayer (Fig. 2C). This indicates that vesicular trafficking of Jam-C is associated with the removal of endothelial cell junctions. We next extended these studies to HUVECs treated with the potent pro-inflammatory cytokines IL-1 and TNF-α. TNF-α increased the number of Jam-C positive intracellular vesicles whilst IL-1 had little effect (Fig. 2D, & E), suggesting that Jam-C junctional remodelling is stimulus specific.

To directly monitor the dynamics of Jam-C trafficking, we transfected HUVECs with a Jam-C-GFPout construct (Fig. 2F) and observed the cells by spinning disk confocal microscopy (Fig. 2G, Movie 1). We noted large membranous structures forming near cellular junctions from which tubular extensions were seen to emerge that subsequently detached and moved away (presumably along cytoskeletal tracks), dissolving the original structure. Together, the present data demonstrates that Jam-C trafficking is a rapid process, a response that is further increased during junctional remodelling and following stimulation of endothelial cells with certain inflammatory stimuli.

### Development of a novel HRP based proximity labelling assay for characterising the trafficking of Jam-C

The endothelial cell junction is comprised of multiple protein complexes, some of which have already been demonstrated to redistribute constitutively and during leukocyte transmigration^36–38^. To define in an unbiased manner the receptors localised with Jam-C at the cell junctions and to determine which of these co-traffic with Jam-C, we developed a Jam-C-HRP based proximity labelling protocol^39^ (Fig. 3A-C). This approach uses the enzymatic activity of HRP to oxidise fluid-phase fed biotin tyramide, thus generating biotin phenoxyl radicals which covalently react with electron rich amino acids (such as Tyr, Trp, His and Cys) on neighbouring proteins (Fig. 3B). The short-lived nature of these radicals results in a small labelling radius (<20nm) and provides a proximity map of all nearby proteins (Fig. 3C). We transfected HUVECs with the Jam-C-HRPout construct (Fig. 3A) and fed the cells for 30 min with biotin tyramide. The proteins neighbouring Jam-C were biotinylated during a short (1 min) incubation with hydrogen peroxide and this reaction was then terminated using molecules that scavenge free radicals. Biotinylated proteins in proximity to Jam-C HRP were detected at the cell surface and within intracellular vesicles (Fig. 3D). Importantly, no biotinylation was detected in the absence of the biotin tyramide or hydrogen peroxide (Supp. Fig. 1A). To specifically identify intracellular stores of Jam-C, we incorporated ascorbate, a membrane impermeant inhibitor of HRP, into our assay. Ascorbate blocks the proximity labelling reaction at the cell surface but not in intracellular stores (Fig. 3C). This approach has been previously utilised in EM studies to inhibit cell surface HRP-catalysed DAB labelling^40, 41^, but to our knowledge this is the first time it has been employed for proteomics. We verified that the reaction worked by immunofluorescent labelling of treated cells with fluorescently-tagged streptavidin to visualise biotinylated proteins near Jam-C (Fig. 3D and Supp. Fig. 1) and by western blotting using streptavidin HRP (Fig. 3E). Both methods confirmed that intracellular stores of Jam-C could be visualised, but not cell surface pools. To further increase the number of intracellular vesicles containing Jam-C, we also incorporated bafilomycin into the assay (Fig. 3D). Following the reaction, cells were lysed and biotinylated proteins pulled down with Streptavidin before performing an on-bead tryptic digest and subsequent tandem mass spectrometry (MS/MS) analysis.

**Figure. 3.**
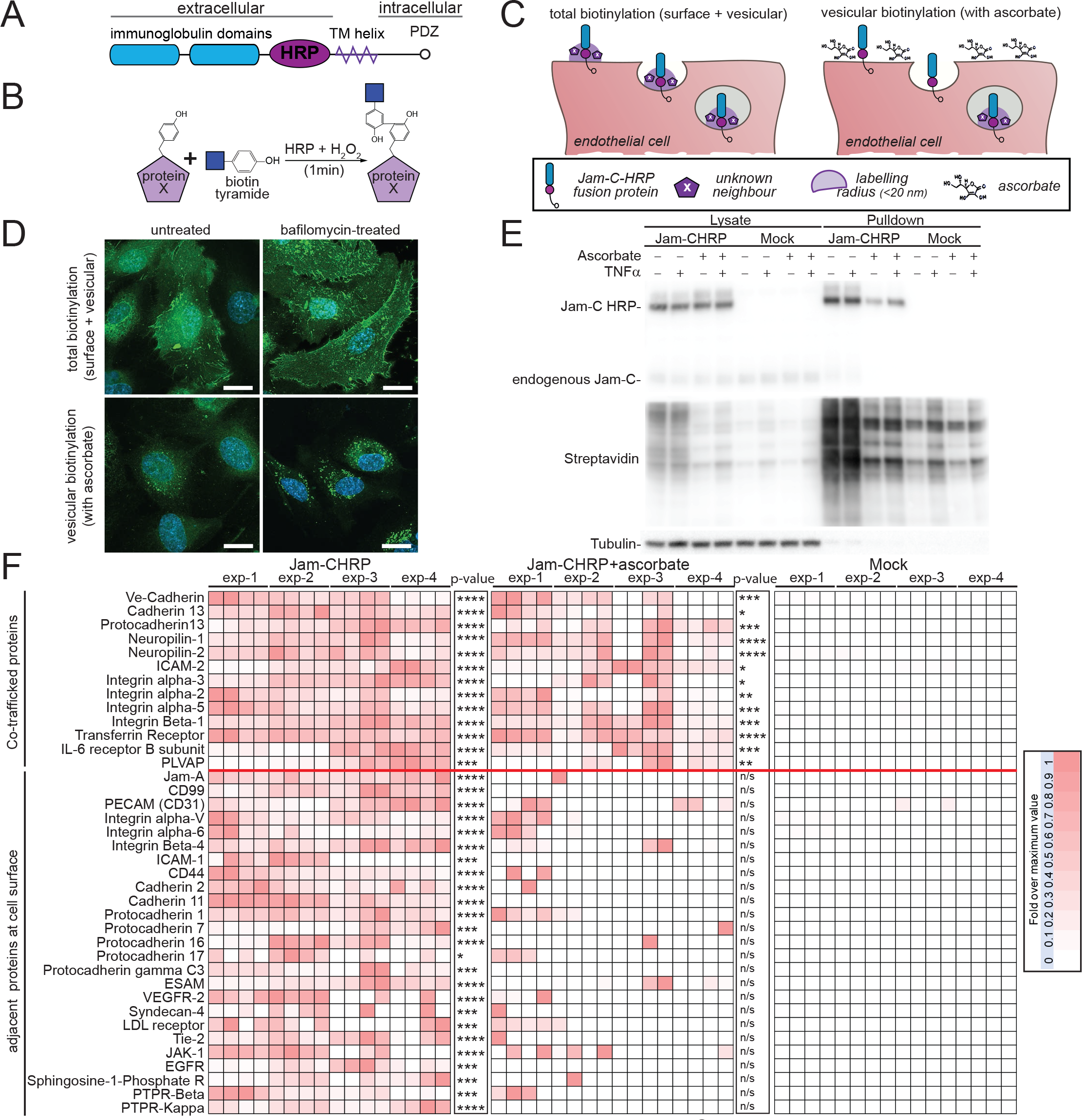
An HRP based proximity labelling approach reveals Jam-C co-traffics with NRP-1, -2 and VE-Cadherin but not with components of the lateral border recycling compartment. (A) Schematic of the domain structure of the Jam-C HRP construct. (B) In the presence of biotin tyramide and hydrogen peroxide the HRP tag leads to the biotinylation of any proteins within 20 nm (labelled with an X) on Tyrosine, Tryptophan, Cysteine or Histidine residues. (C) Endothelial cells expressing Jam-C-HRPout are fed with biotin tyramide and then hydrogen peroxide is added for 1 min. Proteins within 20 nm radius of Jam-C-HRP are labelled with biotin (shown in light purple). Addition of the membrane impermeant HRP inhibitor ascorbate blocks the biotinylation reaction at the cell surface but not inside the cell. (D-F) HUVEC were transfected with Jam-C-HRP out and incubated in the presence or absence of bafilomycin A1 (100 nM) for 4 h to block lysosomal degradation. Cells were fed biotin tyramide for 30 min and then exposed to hydrogen peroxide for 1 min in the presence or absence of 50 mM ascorbate. (D) The cells were then fixed and labelled for streptavidin (green) and DAPI (blue) and images acquired by confocal microscopy. Scale bar 20 μm or (E-F) The reaction was then quenched and the cells lysed. Biotinylated proteins were pulled down using neutravidin beads and lysate and pulldown samples were analysed by (E) SDS PAGE and western blot, for the presence of Jam-C, tubulin and biotinylated proteins or (F) following an on bead tryptic digest mass spectrometry. A heat map of 4 independent mass spectrometry data sets is shown with white none and dark red a high signal. Individual experiments were carried out in duplicate with each mass spectrometry run being repeated twice (to give a total of 4 analyses/experiment). P values are given across all 4 experiments (*P≤0.05, **P≤0.01, ***P≤0.001, ****P≤0.0001 *t*-test). The + TNF experiment is shown in Supp. Fig. 3 Co-trafficked proteins appear in both +/− ascorbate conditions whilst proteins adjacent to Jam-C solely at the cell surface are only present in the – ascorbate condition.

We detected 134 proteins that were within the vicinity of Jam-C at the cell surface and 50 proteins that were proximal to Jam-C after internalisation (Fig. 3F). Our mass spectrometry results (Fig. 3, Supp. Table 3 – includes corrected p-value for multiple comparisons, Supp. 1B&C) indicate that components of a known junctional adhesion molecule trafficking pathway, the lateral border recycling compartment (LBRC) comprising PECAM-1, CD99 and Jam-A^36–38^ are all adjacent to Jam-C at the cell surface but are not co-trafficked with Jam-C. Proteins classically associated with endothelial cell permeability, such as VE-Cadherin, neuropilin-1 and neuropilin-2, were adjacent to Jam-C at the cell surface and also co-trafficked with Jam-C. Other co-trafficked receptors included plasmalemma vesicle associated protein, a protein implicated in permeability, angiogenesis and leukocyte transmigration^42^, and a number of integrin subunits (alpha -2,-3,-5 and beta-1). Notably VEGFR2/KDR, a receptor classically associated with the neuropilins is adjacent to Jam-C at the junctions but absent from the intracellular carriers, suggesting that separate trafficking of these co-receptors may be a means of regulating their functions.

We further validated our MS results by immunofluorescence analysis and western blotting of purified biotinylated proteins (Supp. Fig.2). Since TNFα can increase the number of Jam-C positive vesicles (Fig. 2E), to determine if these are qualitatively different to those in unstimulated cells (i.e. the result of an additional inflammatory pathway), or whether they simply represent an upregulation in the rate of traffic, we carried out HRP proximity labelling in the presence of TNFα. The co-trafficking proteins were largely the same (Supp. Fig. 3). The only major difference noted was a potential reduction in the co-trafficking of Jam-C with VE-Cadherin and this needs to be verified in future work. This indicates that inflammatory stimuli affect the rate but not the primary route of Jam-C intracellular traffic.

### Jam-C turnover is dependent on ubiquitylation of the cytoplasmic tail

To determine the importance of intracellular trafficking on the function of Jam-C we needed to define the mechanism of its turnover. Trafficking of a receptor is most often governed by specific signals and motifs present in the intracellular domain. The cytoplasmic domain of Jam-C is relatively short (44 amino acids) and features a number of conserved potential phosphorylation and ubiquitylation sites as well as a PDZ interacting domain (Fig. 4A). We incorporated mutations of all these sites into our GFPout Jam-C constructs either by site directed or truncation mutagenesis. When expressed in endothelial cells, all mutants exhibited at least some junctional localisation (Fig. 4B). The ΔPDZ mutant exhibited a noticeably different localisation in that it was expressed at higher levels and was also found to be enriched in a peri-Golgi pool. To determine the effect of each mutation on Jam-C-GFP turnover, we transiently transfected HUVECs with the Jam-C constructs and 16 h later added cycloheximide to block new protein synthesis. We then monitored the loss of GFP-labelled protein by western blot over a 24 h period (Fig. 4C and D). We observed that the K283R and the Y267A mutants had slower and quicker turnover than WT Jam-C, respectively. Given that increases in cell surface Jam-C levels are associated with a number of disease states including atherosclerosis and rheumatoid arthritis^19, 24^, and the K283R mutant exhibits a slower turnover, we focused on the potential role of ubiquitylation in Jam-C trafficking and degradation.

**Figure. 4.**
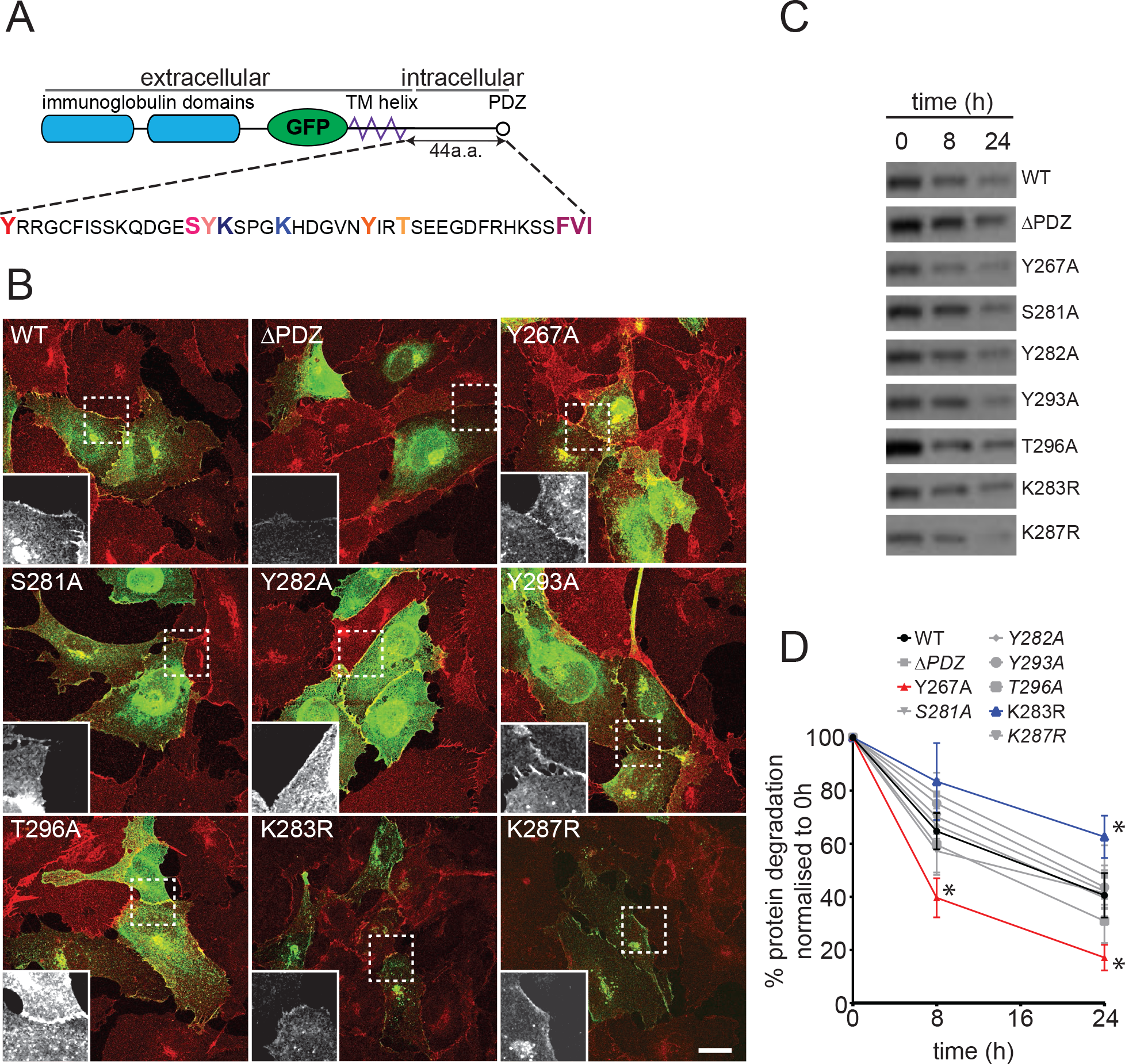
A lysine residue in the cytoplasmic tail of Jam-C is required for its timely degradation. (A) A schematic representation of the Jam-C-GFPout construct showing the residues in the cytoplasmic tail that were mutated by site directed mutagenesis. (B) HUVEC cells were transfected with WT of mutant Jam-C-GFPout constructs, fixed and stained for VE-Cadherin (red). The boxed area is magnified (bottom left of each panel) to show the Jam-C-GFP signal. All Jam-C mutants exhibit at least some junctional staining. The ΔPDZ construct exhibited the most marked change in localisation existing additionally in a peri-Golgi region and expressing at higher levels. Scale bars 20 μm. (C & D) HUVEC were transfected with WT or mutated Jam-C-GFPout constructs and treated with 10 μg/ml cycloheximide. (C) The rate of protein degradation was determined by SDS PAGE and western blot over a 24 h period, a representative anti-GFP blot is shown. (D) Quantification of western blots normalised to the 0 h protein levels, data from n=7 experiments, error bars represent SEM (*P≤0.05; two-way ANOVA).

Jam-C has four lysine residues in its cytoplasmic tail (Fig. 5A); to completely abrogate ubiquitylation, we needed to generate a mutant with all lysine residues mutated to arginine (Quad-K-Jam-C). We co-transfected HUVECs with hemagglutinin (HA)-tagged ubiquitin and either WT or mutant Jam-C-GFPout. Using GFP trap beads, we pulled down the tagged protein and quantified levels of GFP-tagged receptor and associated HA-tagged ubiquitin (Fig. 5B). The Quad-K-Jam-C was devoid of ubiquitylation unlike both the WT and EGFR protein. No obvious change in Jam-C localisation was apparent with the ubiquitin mutant in fixed images (Fig. 5C). However, live cell imaging of WT and Quad-K Jam-C-GFPout in the presence of bafilomycin revealed marked differences in receptor trafficking (Fig. 5D & E). The WT receptor gradually accrues in vesicles following bafilomycin treatment (Fig. 5D, Movie 2). These structures represent the pool of late endosomes and lysosomes unable to degrade due to the inhibition of V-ATPase function and therefore acidification. In contrast, Quad-K Jam-C-GFPout cycles between endocytic structures and the cell surface and does not accrue in late endosomal/lysosomal structures (Fig. 5E, Movie 3). This demonstrates fundamental differences in receptor trafficking following blocking of Jam-C ubiquitylation. To more closely examine the ultrastructure of the intracellular pools in which the Quad-K Jam-C mutant localises, we incorporated the lysine mutations into the Jam-C-HRPout construct and expressed it in unstimulated endothelial cells before carrying out DAB labelling and EM analysis. We found clear evidence of Quad-K Jam-C-HRP on early endosomes (Fig. 5Fa) but little labelling on MVBs (Fig. 5Fb & c), in contrast to WT Jam-C-HRPout (Fig. 1H & I). This data is consistent with the live cell imaging experiments and indicates that the mutation of intracellular lysine residues in Jam-C prevents sorting of the receptor onto the intraluminal vesicles of the MVB for subsequent degradation.

**Figure. 5.**
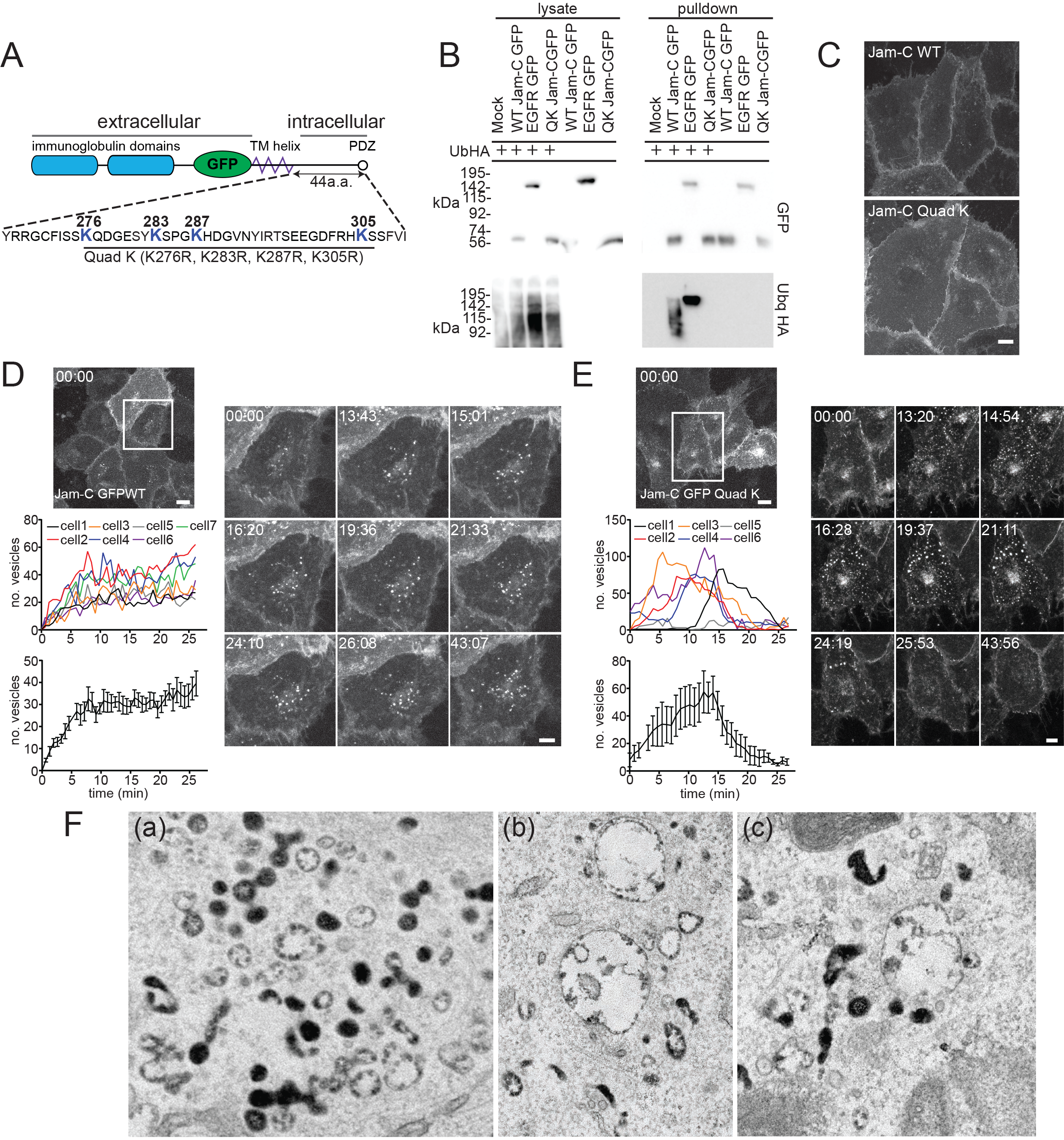
Jam-C is ubiquitylated and this governs its targeting to multi-vesicular bodies. (A) A schematic representation of the Jam-C EGFP Out construct showing all the lysine residues in the cytoplasmic tail that can be potentially modified by ubiquitylation and the Jam-C mutant generated. (B) HUVEC were transfected with WT or Quad K Jam-C-GFPout and Ubiquitin HA. Cells were lysed and GFP tagged proteins were pulled down using GFP trap agarose beads. Lysate and pulldown proteins were analysed by SDS PAGE and western blot for the presence of GFP and HA tagged proteins. WT but not Quad-K Jam-C is modified with ubiquitin. (C) HUVEC were transfected with WT or Quad-K mutant Jam-C-GFPout and then fixed before imaging by confocal microscopy. Scale bar 20 μm. (D & E) HUVEC were transfected with (D) WT or (E) Quad-K Jam-C-EGFPout and 100 nM Bafilomycin was added (at t=0 min) before imaging by timelapse confocal micros-copy. The number of vesicles over time in each cell, or the average number of vesicles/cell are plotted (error bars represent SEM). A representative movie is shown of n=3 separate experiments. Boxed regions are shown magnified as individual stills. Scale bars 20 μm and 5 μm in magnified stills. (F) HUVEC were transiently transfected with Quad-K Jam-C-HRPout, fixed and incubated with diaminobenzidine and hydrogen peroxide for 30 min. Cells were then secondarily fixed and 70 nm sections prepared and imaged by transmission electron microscopy. Precipitated DAB can be clearly seen on early endosomes but there is little evidence of labelling on multi-vesicular bodies (a, b and c) Scale bar 200 nm.

### An APEX-2 proximity labelling approach to identify machinery utilised for Jam-C trafficking

To determine the molecular machinery associated with the intracellular trafficking and ubiquitylation of Jam-C, we used an APEX-2 proximity labelling approach. This strategy was originally developed by Rhee *et al.*^39^. To specifically identify machinery associated with ubiquitylation of Jam-C, we compared WT Jam-C APEX-2 biotinylation with the Quad-K Jam-C APEX-2 and monitored the proteins enriched in the former over the latter.

We incorporated a codon-optimised APEX-2 tag in the membrane proximal region of the cytoplasmic tail of Jam-C in both WT and Quad-K Jam-C (Fig. 6A). This site was chosen so as to minimise the potential for disrupting the C-terminal PDZ interacting domain and other motifs present in the cytoplasmic tail. To ensure the WT and mutant Jam-C APEX constructs did not dimerise with endogenous WT receptor, we depleted the endogenous receptor and expressed siRNA-resistant constructs (Fig. 6B). We detected robust biotinylation by western blot following 30 min feed with biotin tyramide and 1 min exposure to hydrogen peroxide (Fig. 6C). Moreover, biotinylated proteins were detected at the junctions and on intracellular vesicles following immunofluorescence labelling with streptavidin 488 (Fig. 6D). To identify proteins neighbouring Jam-C-APEX, we pulled down biotinylated proteins and carried out on-bead trypsin digestion and MS/MS analysis. MS analysis revealed 576 significant hits that were more than 3.5 fold enriched over the mock transfected cells (Fig. 6E, F, & Supp. Table 4). Hits included proteins from the cell surface and intracellular pools and associated with a number of cellular processes including endocytosis, recycling, fusion and ubiquitylation. We detected a number of integral membrane proteins previously identified from our HRP approach including VE-Cadherin. Of particular interest were a number of components associated with ESCRT-0,-1 and E3 ligases such as CBL, demonstrating that at some point during its trafficking, Jam-C is adjacent to machinery necessary for the ubiquitylation and the maturation of endosomes to MVBs. The equivalent analysis of Quad-K Jam-C-APEX-2 biotinylation showed a similar profile but, notably, CBL, and components of the ESCRT complex VPS28 and Vps4B were absent (Fig. 6E & G). This indicates that, when the 4 lysines present in the cytoplasmic tail of Jam-C are mutated, the E3 ligase and some components of the associated ESCRT machinery are no longer in proximity to Jam-C (the lack of ubiquitylation sites likely reduce the amount of time CBL localises to the receptor). The absence of ubiquitylation results in a failure to sort Jam-C onto intraluminal vesicles.

**Figure. 6.**
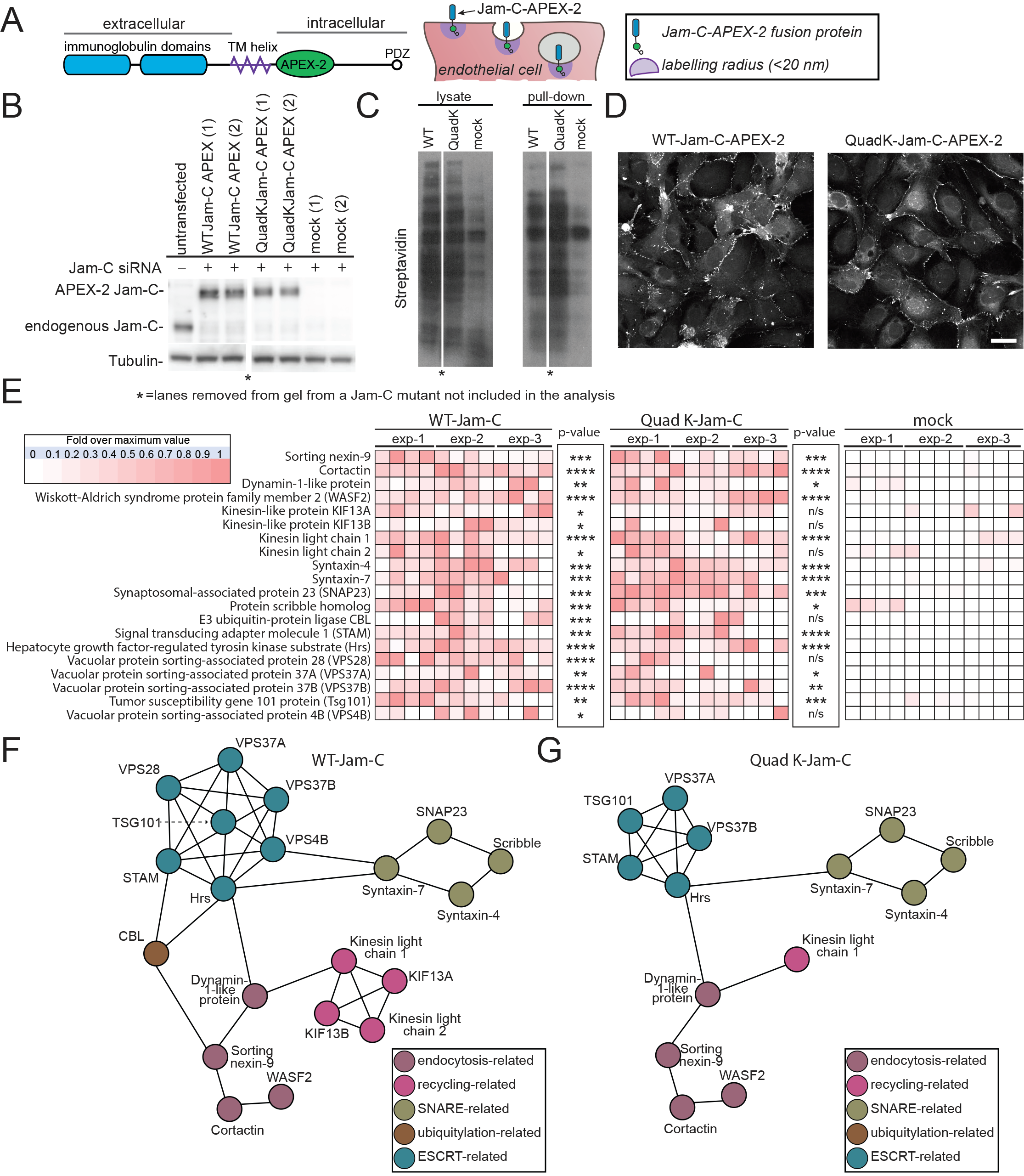
An APEX-2 proximity labelling approach reveals potential Jam-C trafficking machinery. (A) Schematic of the domain structure of the Jam-C-HRP-APEX-2in construct showing the extracellular immunoglobulin domains (blue), the APEX-2 tag (green oval) and the intracellular PDZ interacting domain (clear circle). The APEX-2 tag following feeding with biotin tyramide and incubation with hydrogen peroxide for 1 min labels all neighbouring proteins within 20 nm on electron rich amino acids (purple circle). (B-E) HUVEC were transfected with WT or Quad-K Jam-C-APEX-2in. Cells were fed biotin tyramide for 30 min and then exposed to hydrogen peroxide for 1 min. The reaction was then quenched and the cells either (D) fixed and stained with streptavidin and analysed by confocal microscopy (B,C & E) lysed and biotinylated proteins pulled down using neutravidin beads for analysis by (B & C) SDS PAGE and western blot for the presence of (B) Jam-C and tubulin or (C) biotinylated proteins or (E) following an on bead tryptic digest, mass spectrometry. A heat map of n=3 independent mass spectrometry data sets is shown with white representing no signal and dark red a high signal. Each individual experiment was carried out in duplicate with each mass spectrometry run being repeated twice (to give a total of 4 analyses/experiment). P values are given across all 3 experiments (*P≤0.05, **P≤ 0.01, ***P≤0.001, ****P≤0.0001; t test). (F & G) String analysis of proteins neighbouring (F) WT or (G) Quad-K Jam-C APEX-2 proteins. *lane removed of a Jam-C mutant not included in this analysis.

To confirm that CBL is required for the ubiquitylation of Jam-C we reduced its expression using siRNA transfection in HUVECs (Fig. 7A). Depletion of CBL decreased the ubiquitylation of Jam-C-GFPout (Fig. 7B & C) and increased the levels of receptor present in endothelial cells (Fig. 7B and D). The effects of CBL depletion are particularly apparent when the levels of ubiquitylation are normalised to the levels of Jam-C-GFPout present (Fig. 7E). CBL knockdown caused no obvious change in endogenous receptor localisation by immunofluorescence analysis (Fig. 7F). However, on addition of bafilomycin, CBL depleted cells have significantly fewer Jam-C-GFPout positive vesicles (Fig. 7F & G). This indicates that CBL is required for Jam-C ubiquitylation, which is necessary for Jam-C sorting onto the intraluminal vesicles of late endosomes.

**Figure. 7.**
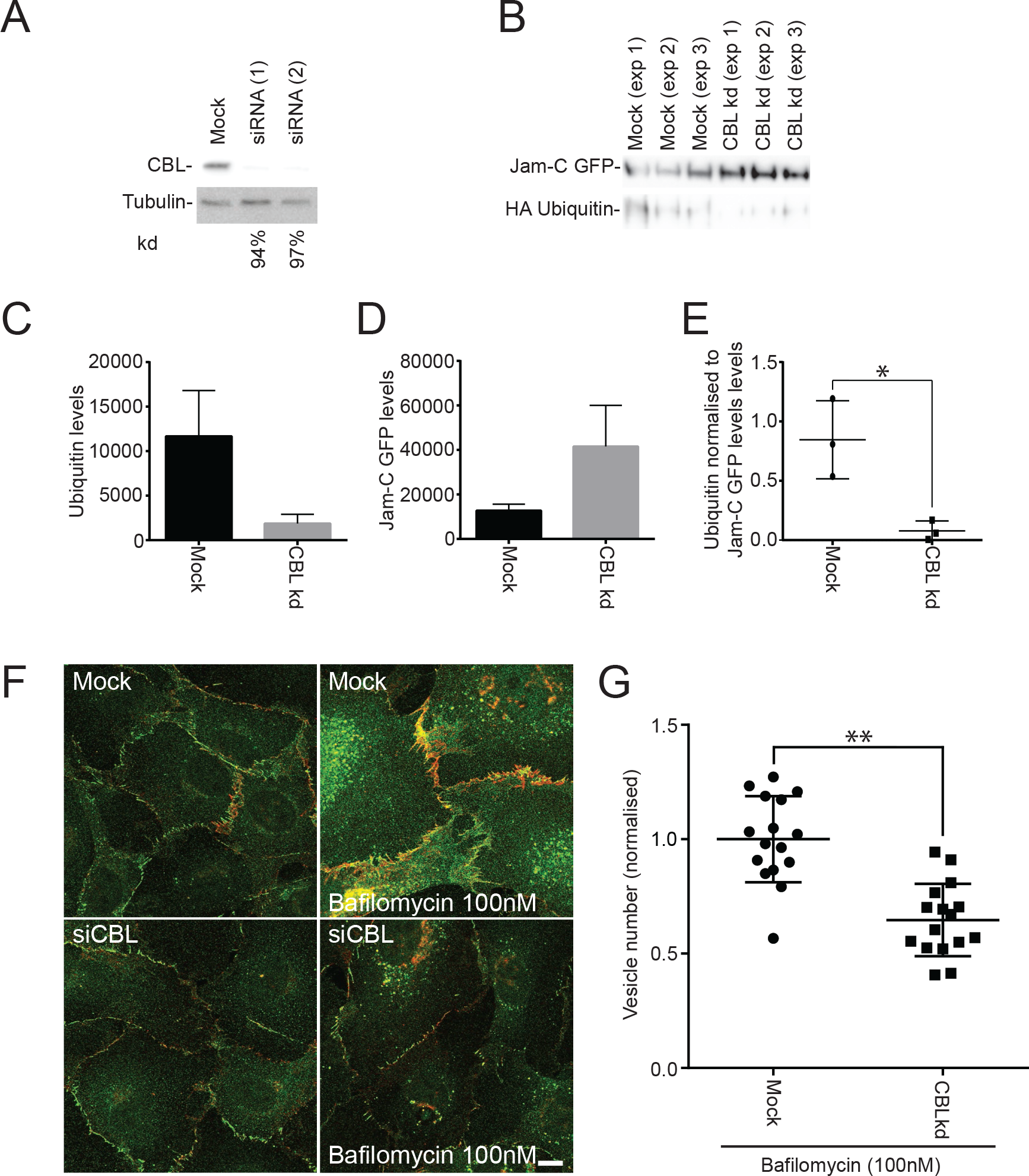
Jam-C is ubiquitylated by the E3-ligase CBL. (A-G) HUVEC were transfected with two different CBL siRNA over a 96 h period. (A) Cells were lysed and analysed by SDS PAGE and western blot detecting CBL and tubulin. Knock down was typically ≥ 90%. (B) Untreated or CBL knock down cells were transfected with Jam-C-GFPout and HA ubiquitin. Cells were lysed and GFP tagged proteins were pulled down using GFP trap agarose beads. (C-E) Pulldown proteins were analysed by SDS PAGE and western blotted for the presence of GFP and HA tagged proteins. Quantification of (C) HA ubiquitin levels (D) Jam-C-GFPout levels or (E) Ubiquitin levels normalised to levels of Jam-C. Results are from n=3 experiments. Error bars represent SEM (*P≤0.05; t test). (F) Mock and CBL knockdown cells were incubated with and without 100nM Bafilomycin for 4 h and then fixed and labelled for endogenous VE-Cadherin (red) and Jam-C (green). (G) Quantification of the number of Jam-C positive vesicles present in mock and CBL knockdown cells treated with 100 nM bafilomycin (5 fields of view/condition). Results are shown normalised to the mean number of vesicles in the mock cells from n=3 experiments. Error bars represent SD (**P≤0.01; t test).

### Ubiquitylation of Jam-C is required for cell migration

Jam-C is known to mediate a number of important biological processes including angiogenesis^16, 43-46^ and leukocyte transmigration^8–12^. To determine how altering the ubiquitin mediated-trafficking of Jam-C affects its cellular function, we performed scratch wound assays. Recovery of the wound has been shown to be ameliorated in conditions of Jam-C knock down^47^ and increased in situations of Jam-C overexpression^48^. This assay provides a simple way to monitor the migration of endothelial cells, a process which is essential during angiogenesis. Such changes are unlikely to be due to altered proliferation as no change in cell growth has been detected following Jam-C over-expression over a 96 h period^49^. We depleted endogenous Jam-C and rescued with either lentiviral expressed GFP, WT Jam-C-GFPout or Quad-K Jam-C-GFPout (Fig. 8A) and monitored the rate of scratch wound closure over a 16 h period (Fig. 8B). The efficiency of endogenous Jam-C knockdown was similar in all conditions, and the expression levels of the Jam-C constructs matched endogenous levels (Fig. 8A). Knockdown of Jam-C resulted in a slower rate of scratch wound closure and this was rescued by overexpression of Jam-C-GFPout (Fig. 8B & C). In contrast, Quad K Jam-C-GFPout failed to rescue the phenotype (Fig. 8B & C) and 60 h post scratch the wound remained open (Fig. 8D).

**Figure. 8.**
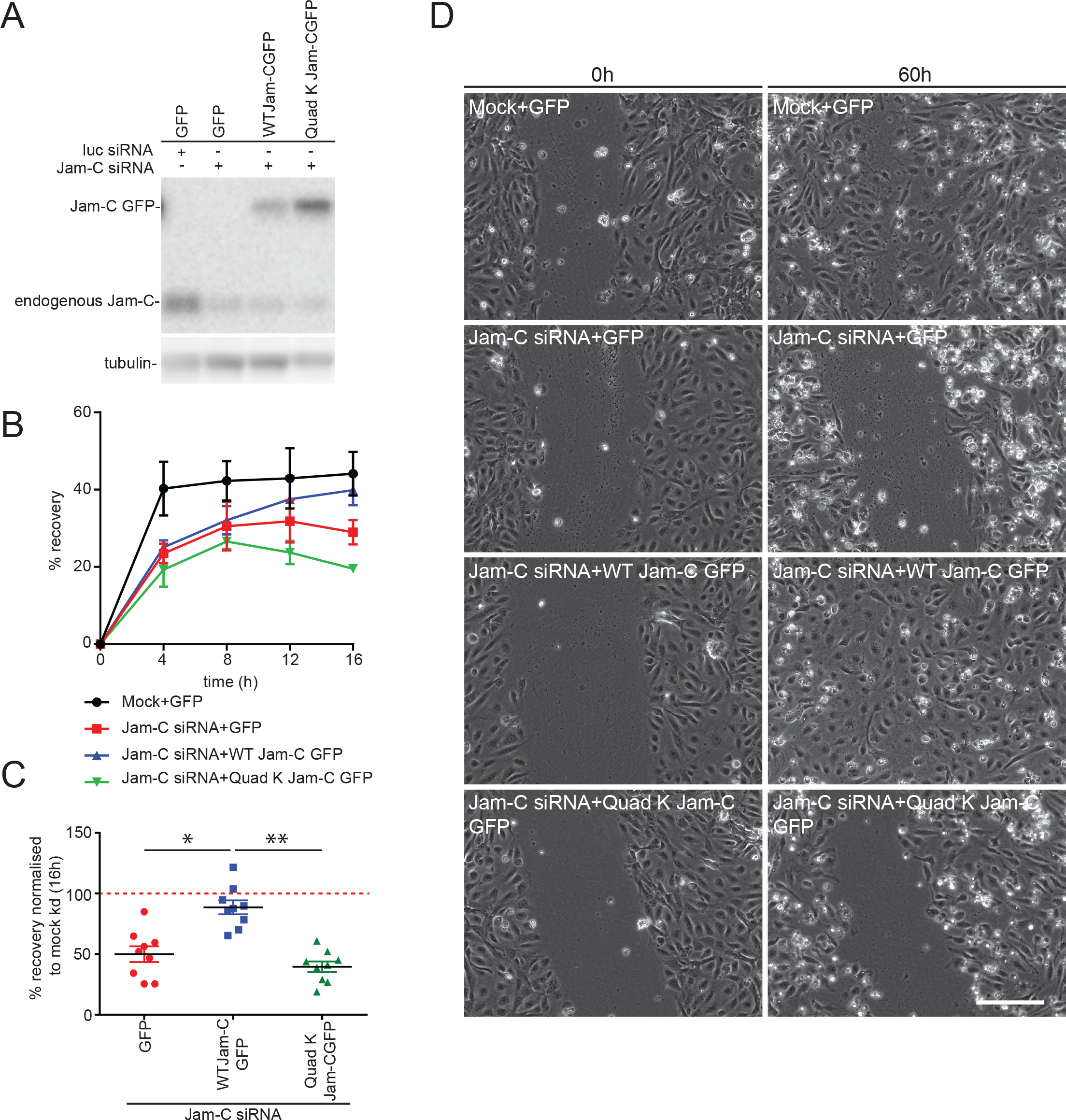
Ubiquitylation of Jam-C is required for HUVEC migration. (A-D) HUVEC were transfected with siRNA targeting luciferase or Jam-C over a 96h period. The day after the second round of transfection cells were additionally transduced with either a GFP, WT Jam-C-GFPout or a Quad K Jam-C-GFPout expressing lentivirus and allowed to reach confluency. (A) Cells were lysed and the extent of Jam-C knock down and rescue determined by western blot relative to a tubulin loading control. The expressed WT and Quad K constructs were similar to endogenous levels. (B-D) The monolayer was scratched and the cells allowed to recover over a 0-60 h time period whilst being imaged on a brightfield Olympus microscope. (B) A representative experiment showing the percentage recovery of cells over a 16 h time period. Mock treated cells expressing the GFP construct recover relatively quickly. Jam-C siRNA treated cells fail to recover when rescued by transduced QuadK Jam-C-GFPout or GFP construct. However, almost complete recovery is apparent using transduced WT Jam-C-GFPout construct after 16 h. Error bars represent standard deviation. (C) The percentage recovery of siRNA treated GFP, WT Jam-CGFP or QuadK Jam-CGFP transduced cells at 16 h is shown normalised to the mock/GFP rescued condition. Data shown from n=4 experiments, error bars represent SEM (*P≤0.05, ** P≤0.01; unpaired t-test). (D) Representative images of recovery of scratch wounds after 60 h. Scale bar 200 μm.

Finally, we used live cell imaging to directly analyse the trafficking of Jam-C in migrating cells (Fig. 9A, Movie 4). We noted large Jam-C-GFPout positive vesicles forming at junctional sites on either side of the cell’s leading edge; no such vesicle traffic was apparent at the very front of migrating cells. This indicates that junctional disassembly is a crucial aspect of cell migration and a proportion of this internalised Jam-C needs to be degraded by a ubiquitin-mediated lysosomal pathway to allow normal cell migration (Fig. 8). Together, the results demonstrate dynamic Jam-C trafficking and ubiquitylation are essential responses for endothelial cell migration.

**Figure. 9.**
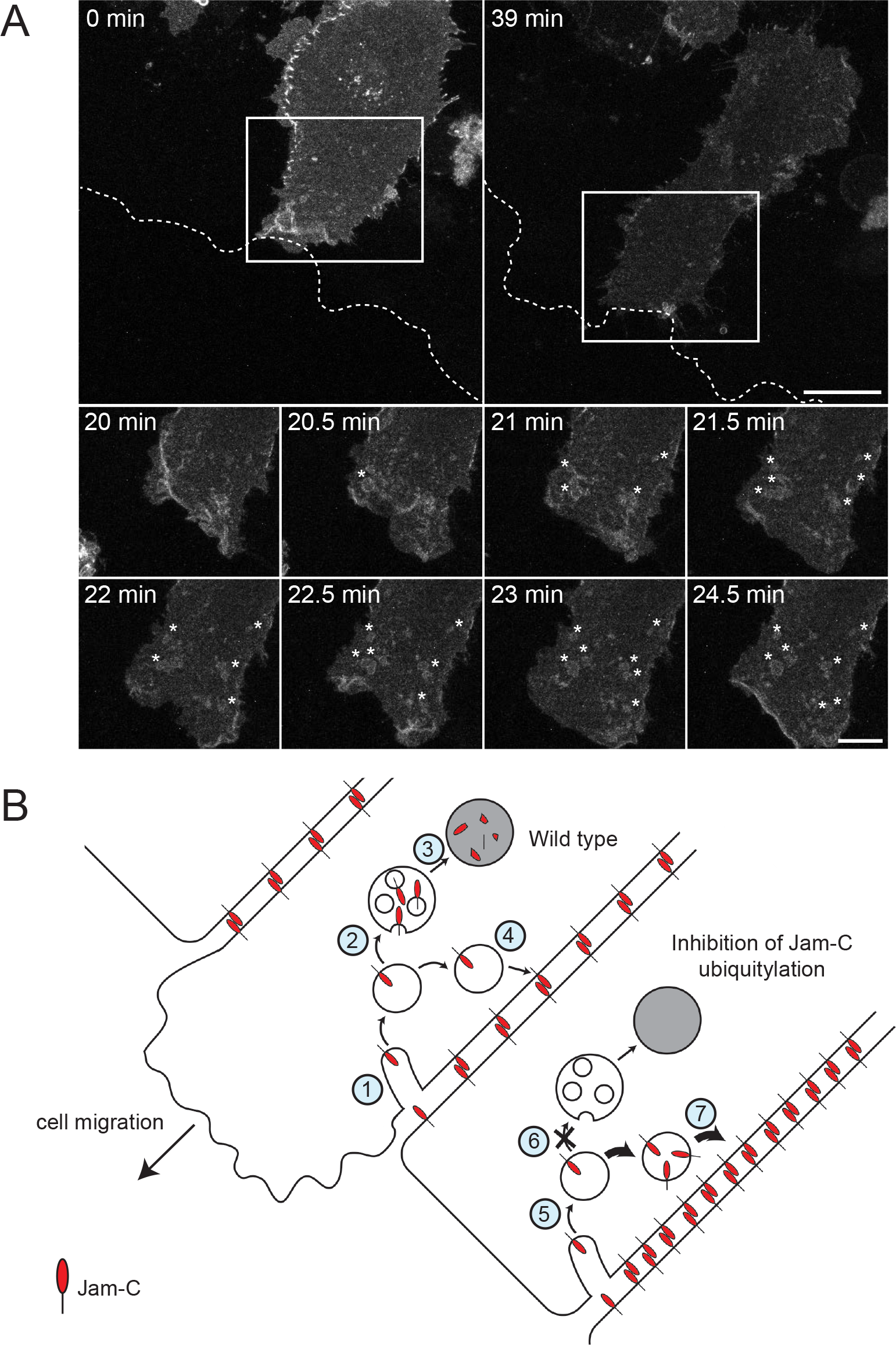
Jam-C is internalised from junctions near the front of migrating cells. (A) Jam-C-GFPout was expressed in HUVEC and the cells allowed to form a confluent monolayer. The cell layer was scratched and 45 min later live cell imaging was carried out with images acquired every 30 s for 40 min. The dotted line represents the edge of the scratch wound (as determined by DIC imaging), asterisks highlight vesicles and the boxed region is shown magnified. Scale bar 20 μm in original images and 10 μm in magnified images. (B) Model of the role of Jam-C trafficking in endothelial cell migration. **Wild type**: (1) Jam-C is endocytosed from the junction either side of the leading edge and is localised to endosomes. (2) As Jam-C enters endosomes the E3 ligase CBL ubiquitylates the cytoplasmic tail on lysine residues, allowing recruitment of the ESCRT complex and the formation of intra-luminal vesicles. (3) Jam-C is degraded in the lysosomes preventing excessive recycling and/or abrogating signalling. (4) The remaining Jam-C that is not ubiquitylated is recycled to the cell surface to reform new junctions after the cell has moved. **Inhibition of Jam-C ubiquitylation**: (5) Jam-C is endocytosed from the junction either side of the leading edge and is localised to endosomes. (6) Mutation of Jam-C or absence of CBL prevents ubiquitylation and recruitment to the intraluminal vesicles of late endosomes. (7) Failure to degrade Jam-C results in an increased premature recycling of Jam-C to the cell junction resulting in greater adhesion between cells and/or enhanced signalling.

## Discussion

Prior to this study many aspects of the molecular control of Jam-C function were unclear. As was the relationship, in terms of regulation and function, with its homologue Jam-A and other components of the junctional machinery. Our presented work addresses many of these issues and opens further avenues of research to determine how intracellular trafficking controls Jam-C’s important physiological and pathological roles.

Our proximity labelling proteomics approach allowed a bulk analysis of the receptors trafficked alongside Jam-C. With this approach we discovered that Jam-C traffics with VE-Cadherin, NRP-1 and -2 as well as some integrin subunits, but is not present in the same pool as its orthologue Jam-A or any of the other component of the LBRC. This co-trafficking is likely to represent receptors associated with a similar function being moved en masse. Receptors present in the LBRC are thought to provide a reticular pool of unligated receptors necessary for leukocyte movement whilst by contrast we hypothesise trafficking of Jam-C and its counterparts allow precise control of cell migration and the passage of molecules and cells by disassembling and reforming the junction. In agreement with this we see increased trafficking of Jam-C associated with artificial disruption of the junctions by calcium chelation and following exposure to an inflammatory stimulus (TNFα). Another cytokine, IL-1 had little effect on trafficking and this difference likely reflects the ability of TNFα signalling to additionally stimulate an endothelial permeability response^50^. *In vivo* TNFα signalling is known to be important during ischemia reperfusion injury^51^ one of the first occasions in which vesicular pools of Jam-C were noted^10^.

Although there is no direct co-dependence for traffic, it remains possible that Jam-C may share some as yet unknown trafficking machinery with its neighbours and be internalised at the same time from the cell surface. However, some of the receptors present in the same co-trafficked pool have been reported to be internalised by different clathrin dependent and independent routes ^52–56^. An alternative explanation is that the shared pool could represent a fusion of early endosomes from diverse endocytic routes to form a late endosomal pool downstream. There is therefore scope for co-trafficked receptors interacting with each other and therefore modifying function although this remains to be investigated.

A key finding of the study is that we show for the first time that Jam-C is ubiquitylated and that this modification represents a key step in Jam-C receptor trafficking. Our APEX-2 proximity labelling approach allowed us to identify CBL as the E-3 ligase responsible for ubiquitylating Jam-C on up to 4 different residues in the cytoplasmic domain, with the number and the pattern of bands suggesting polyubiquitylation. This modification is necessary for the efficient sorting of Jam-C onto intraluminal vesicles of the MVB. The trafficking phenotype on CBL depletion was less striking than that of the Quad-K Jam-C mutant, and so likely represents either an incomplete block of the ubiquitylation pathway or compensation by another E3 ligase. Ubiquitylation is a means of controlling turnover of claudins^57^ however the only Jam family member previously shown to be ubiquitylated is Jam-B on Sertoli cells, the site of ubiquitylation, the E3 ligase involved, and the effect on receptor trafficking has yet to be determined^58^. A further means to regulate Jam-C trafficking and function is likely to be the removal of ubiquitin. There are thought to be more than 100 potential deubiquitinating enzymes (DUB) in mammalian cells^57^ and we identified 6 potential Jam-C localised DUBs in our APEX-2 screen USP-5, -7, -10, -15, -47 and USPY. Notably, these proteins were adjacent to Jam-C irrespective of whether we utilised the WT or the Quad-K mutant and therefore could potentially represent nearby bystander proteins rather than bona fide interactors. We have yet to verify if these proteins are important for Jam-C traffic although interestingly USP-47 has been shown to regulate E-cadherin deubiquitylation at junctions^59^.

Our results show that targeted removal and degradation of Jam-C from junctions is required for endothelial cell migration. As endothelial cells migrate to fill a gap in the monolayer, endocytic disassembly of the junction occurs either side of the cells leading edge. As Jam-C enters endosomes it becomes ubiquitylated and is directed for degradation by the lysosomes. A failure to ubiquitylate has no effect on internalisation but results in premature recycling of Jam-C back to the cell surface. We present two potential models (which are not mutually exclusive) of how inhibition of degradation of Jam-C slows migration (Fig. 9B). Firstly, the change in receptor recycling might prematurely increase the strength of homo and heterotypic junctional interactions with neighbouring cells preventing cell movement. Secondly, targeting receptor for degradation might be required to disrupt cell surface protein-protein interactions or terminate signalling events. Such signalling could potentially occur via interactions with the polarity proteins Par3^60^ and Par6^61^. Notably Par3/Jam-C interactions are required for neuronal cell migration from the germinal zone^62^ and signalling via Par3 and Cdc42 is required for lumen formation during 3D endothelial tubulogenesis^63^.

The trafficking and degradation of Jam-C is likely important for the other ascribed roles of Jam-C, particularly during leukocyte transmigration and permeability as they both rely on the disassembly and reassembly of junctions. Controlling the trafficking of Jam-C and its co-trafficked counterparts could potentially provide novel ways of limiting disease states, particularly those associated with increased levels of cell surface Jam-C, such as atherosclerosis and arthritis.

## Acknowledgments

T.D. Nightingale, C. Schultz and K. Kostelnik were funded by an MRC project grant MR/M019179/1. The research leading to these results has also received funding from the People Programme (Marie Curie Actions) of the European Union's Seventh Framework Programme (FP7/2007-2013) under REA grant agreement n° 608765. A. Barker was funded by QMUL. S. Nourshargh was funded by a Wellcome Trust investigator award 098291/Z/12/Z. M. Aurrand Lions was funded by Canceropôle PACA (Valo-Paca 2016) and French National Institute of Cancer (Inca, PRT-K16, #2017-24). P. Cutillas and V. Rajeeve were funded by BBSRC (BB/M006174/1) and the Barts and The London Charity (297/2249). I.J. White was funded by an MRC LMCB core grant award MC_U12266B.

---

**Supplementary Figure. 1.**
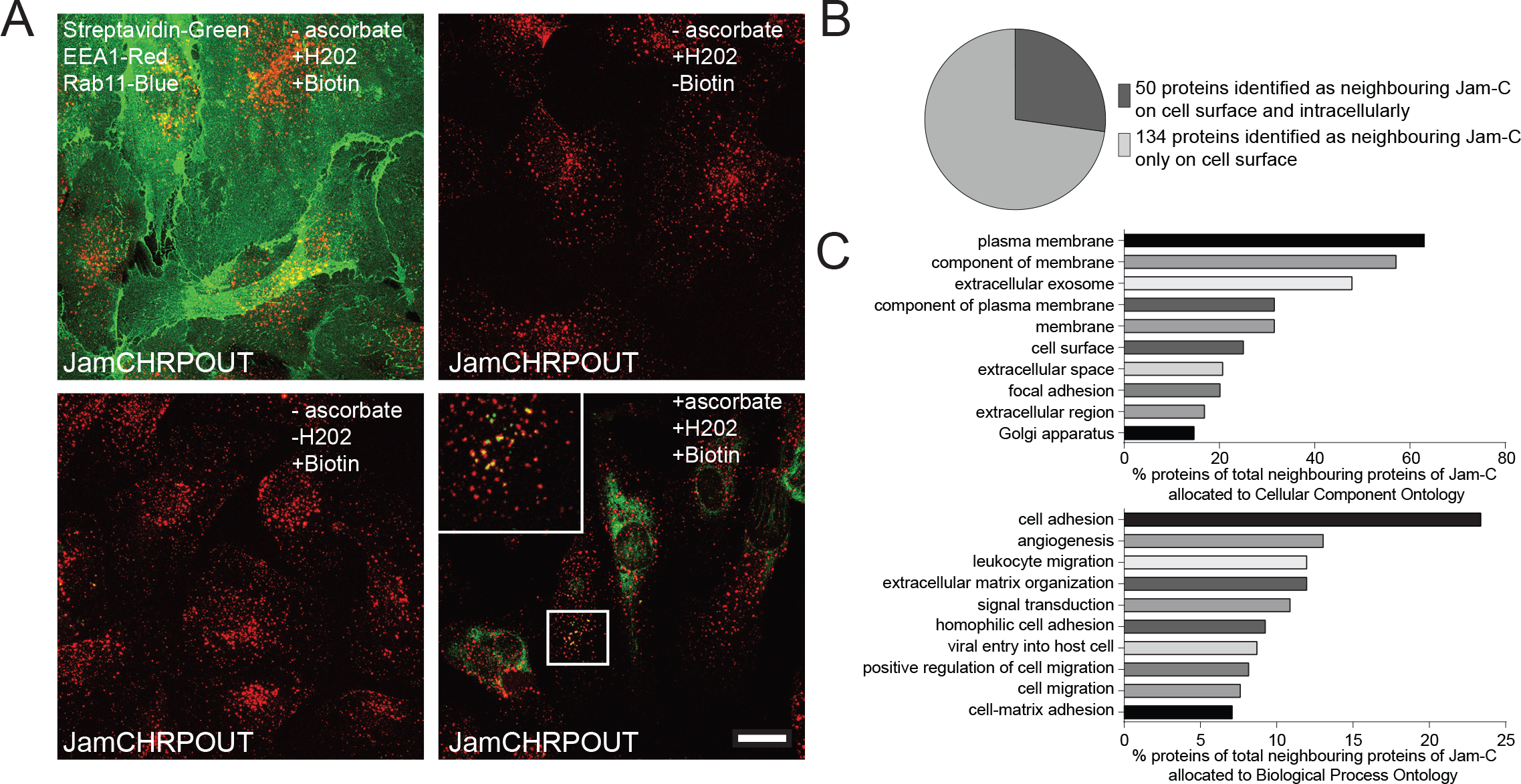
The Jam-C HRP biotinylation assay. (A-C) HUVEC were transfected with Jam-C HRP and the following day incubated with or without biotin tyramide and hydrogen peroxide in the presence or absence of 50 mM ascorbate. (A) Immunofluorescence analysis of cells labelled for Streptavidin (Green) and early endosome antigen 1(EEA 1, Red). Biotinylated proteins are only present when both hydrogen peroxide and biotin tyramide is present and partially localise with early endosomes. (B-C) Biotinylated proteins were pulled down with streptavidin and analysed by mass spectrometry (n=4 experiments) (B) Pie chart showing the number of proteins adjacent to Jam-C at the cell surface and intracellularly. (C) The percentage of protein hits associated with specific cellular locations and processes is plotted.

**Supplementary Figure. 2.**
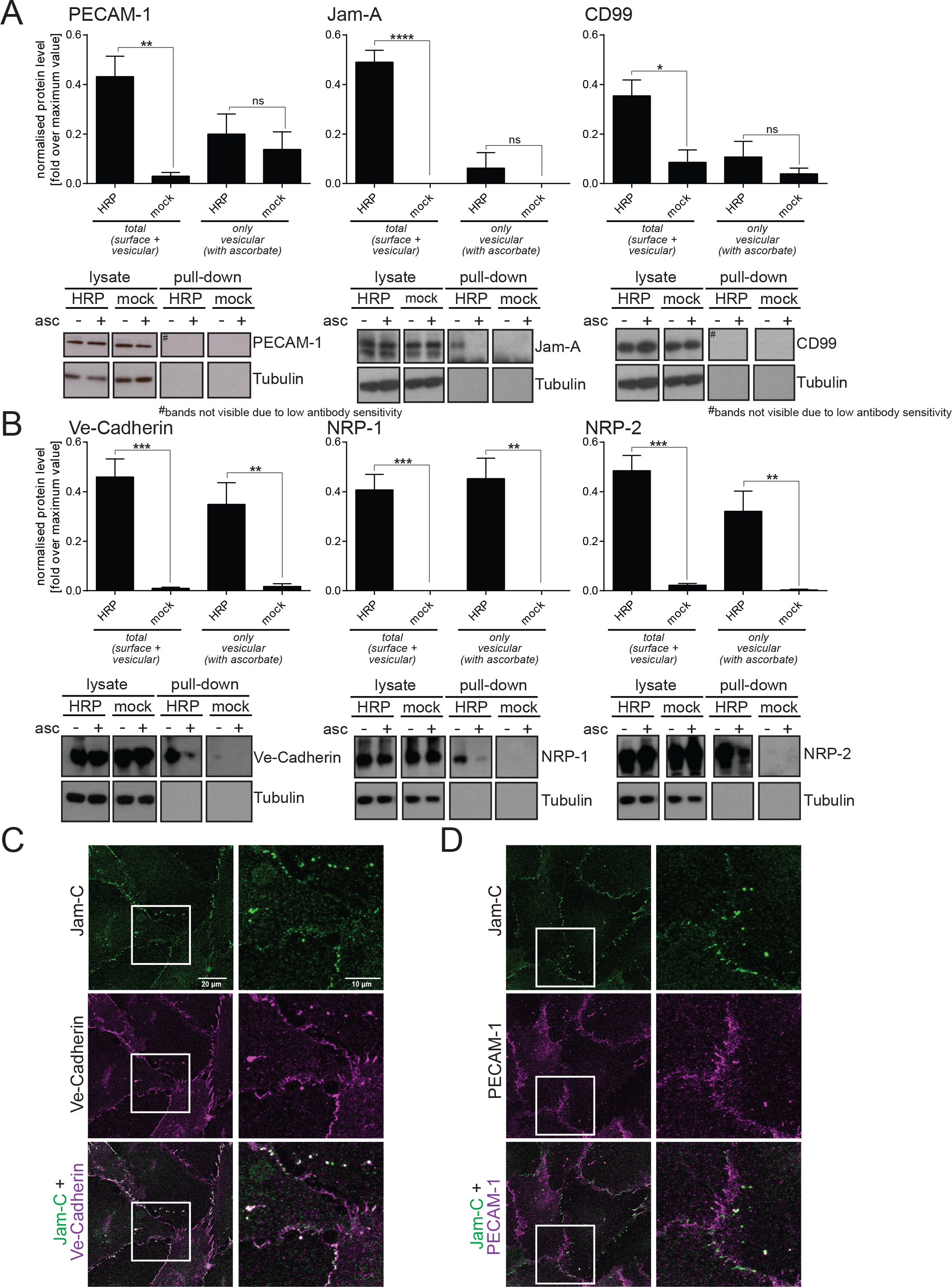
Validation of HRP biotinylation assay by western blot and Immunofluorescence analysis. (A & B) Jam-C-HRPout transfected cells were fed with biotin tyramide and exposed to hydrogen peroxide in the presence or absence of ascorbate. Biotinylated proteins were pulled down and western blotted for (A) proteins neighbouring Jam-C at the cell surface: PECAM-1, Jam-A and CD99 or (B) proteins co-trafficked with Jam-C: VE-Cadherin, NRP-1 and NRP-2. Representative blots are shown with quantification of n=4 experiments, error bars represent SEM (*P≤0.05, ** P≤0.01; ***P≤0.001, ****P≤0.0001; unpaired t-test). (C) Immunofluorescence analysis of endogenous Jam-C (green) and either VE-Cadherin or PECAM-1 (magenta). The boxed region is magnified. VE-Cadherin co-traffics with Jam-C whilst PECAM-1 does not.

**Supplementary Figure. 3.**
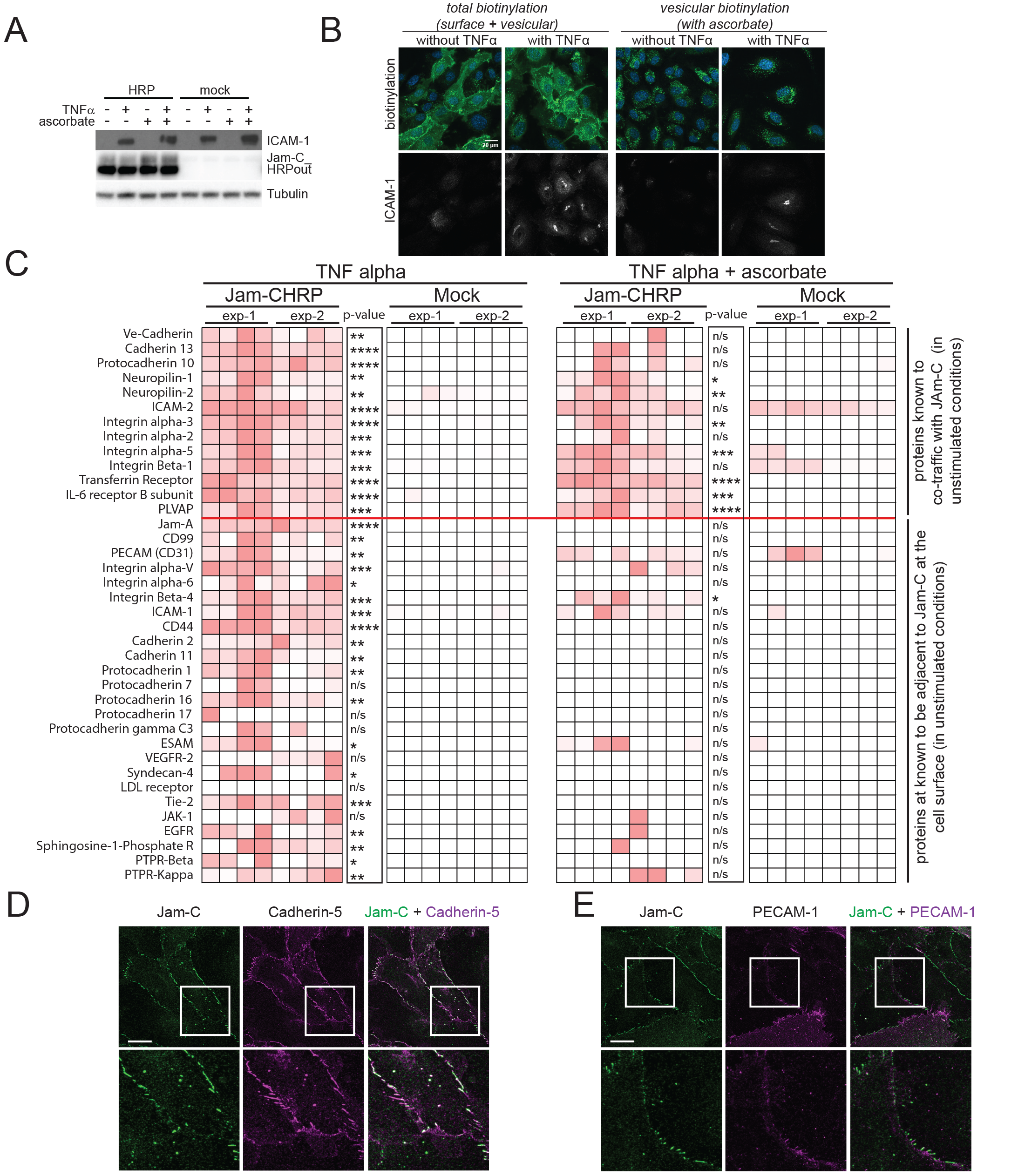
An HRP based proximity labelling approach reveals changes in Jam-C co-trafficking following stimulation with TNFα. (A-C) HUVEC were transfected with Jam-CHRP out and stimulated for 4 h with 50 ng/ml TNFα. (A) Cells were lysed and analysed by western blot. The level of Jam-C HRP expression is similar across all transfected samples and TNFα stimulation upregulates the expression of ICAM-1. (B & C) Cells were fed biotin tyramide for 30min and then exposed to hydrogen peroxide for 1min in the presence or absence of 50mM ascorbate. (B) Cells fixed and stained with streptavidin (green), DAPI (blue) and ICAM-1 (grey). Images were acquired by confocal microscopy. Scale bar 20 μm. (C) Biotinylated proteins were pulled down using neutravidin beads and pulldown samples were analysed by mass spectrometry. Heat map of 2 independent mass spectrometry data sets is shown with white no signal and dark red a high signal. Each individual experiment was carried out in duplicate with mass spectrometry runs being repeated twice (to give a total of 4 analyses/experiment). P values are given across 2 experiments (*P≤0.05, **P≤0.01, ***P≤0.001, ****P≤0.0001; t-test). Co-trafficked proteins appear in both +/− ascorbate conditions whilst proteins adjacent to Jam-C solely at the cell surface are only present in the – ascorbate condition. (D & E) HUVEC were stimulated for 4 h with TNFα fixed and labelled for (D) Jam-C (Green) and VE-Cadherin (magenta) or (E) Jam-C (Green) and PECAM-1 (Magenta). Jam-C does not co-localise with VE-Cadherin or PECAM-1. Scale bar 20 μm.

**Supplemental Table 1.**
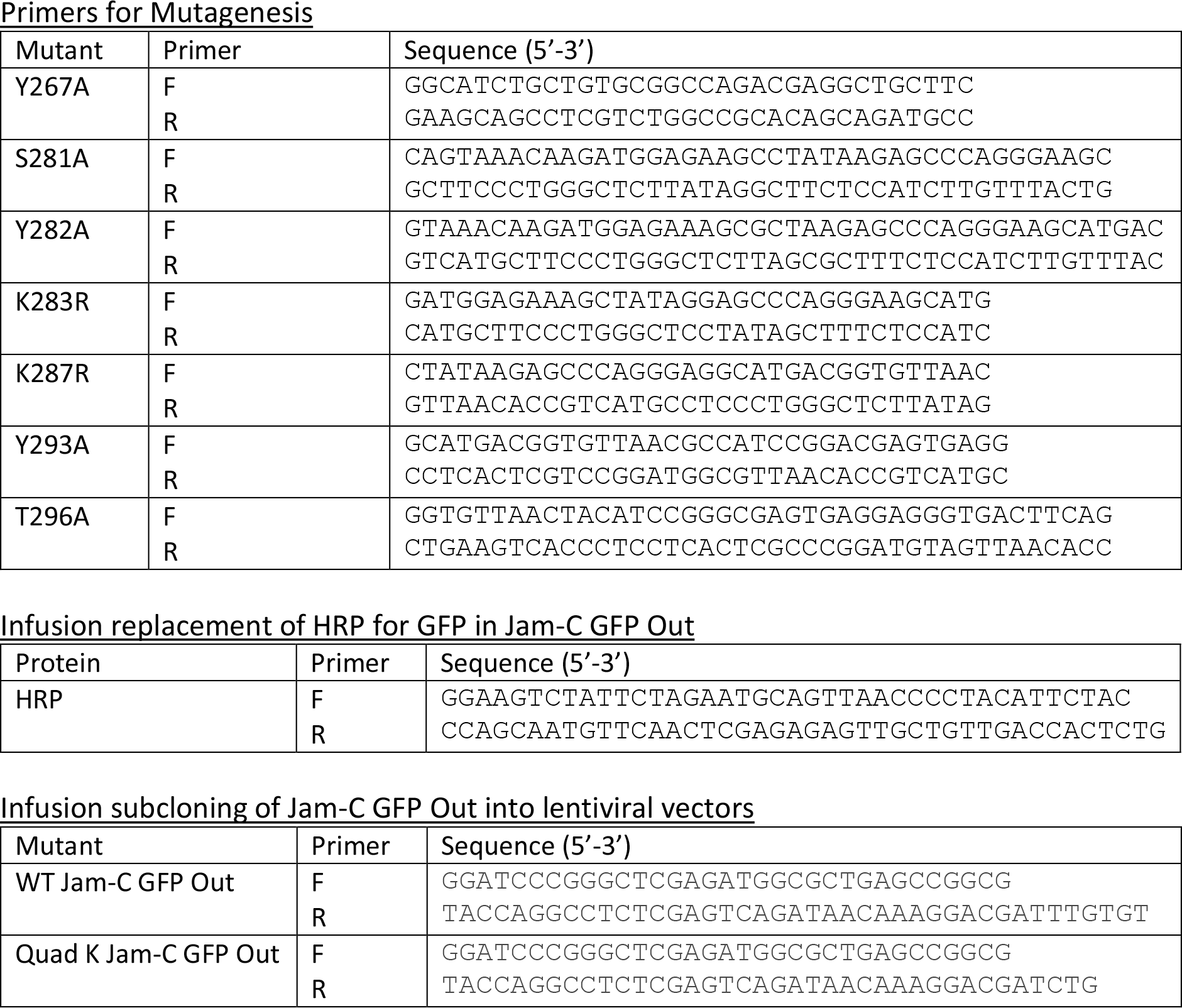
Primer Sequences.

**Supplementary Table 2.** Antibodies.

| Antibody | Source | Cat #/Ref | Species | Details |
| --- | --- | --- | --- | --- |
| HA | Roche | 118674230<br>01 | Rat monoclonal | Western 1:250 |
| GFP | Chromotek | 3H9 | Rat monoclonal | Western 1:1000 |
| Jam-C | A kind gift from Prof Beat Imhof (University of Geneva) | J81 | Western 1:1000<br>IF 1:1000 | Western 1:1000<br>IF 1:1000 |
| Jam-C | Bethyl Laboratories | A303-761A | Rabbit polyclonal | Western 1:1000 |
| VE-cadherin | Santa Cruz | Sc9989 | Mouse monoclonal | Western 1:1000<br>IF 1:100 |
| ZO-1 | Cell signalling technology | D6L1E | Rabbit monoclonal | Western 1:1000<br>IF 1:100 |
| CD63 | AbCAM | CLB180 | Mouse monoclonal | IF 1:100 |
| Rab11 | Lifetechnologies | 71-5300 | Rabbit polyclonal | IF 1:100 |
| NRP-1 | AbCAM | Ab81321 | Rabbit polyclonal | Western 1:1000<br>IF 1:100 |
| NRP-2 | R & D Signalling | AF2215 | Rabbit polyclonal | Western 1:200<br>IF 1:50 |
| PECAM (CD31) | E-bioscience | 17-0319-41 | Mouse monoclonal (WM-59) | IF 1:200 |
| PECAM (CD31) | Santa Cruz | Sc1506 | Goat polyclonal | Western 1:1000 |
| Jam-A | Santa Cruz | Sc53628 | Mouse monoclonal | Western 1:1000<br>IF 1:100 |
| CD99 | Santa Cruz | Sc28389 | Mouse monoclonal (12E7) | Western 1:100 |
| Tubulin | Sigma | T4026 | Mouse monoclonal (Tub2.1) | Western 1:1000 |
| Streptavidin HRP | DAKO | P0397 | - | Western 1:1000 |
| ICAM-1 | Santa Cruz | Sc7891 | Rabbit polyclonal | Western 1:1000 |
| CBL | Santa Cruz | Sc398282 | Mouse monoclonal | Western 1:1000 |

